# Phylogenomic analyses of non-Dikarya fungi supports horizontal gene transfer driving diversification of secondary metabolism in the amphibian gastrointestinal symbiont, *Basidiobolus*

**DOI:** 10.1101/2020.04.08.030916

**Authors:** Javier F. Tabima, Ian A. Trautman, Ying Chang, Yan Wang, Stephen Mondo, Alan Kuo, Asaf Salamov, Igor V. Grigoriev, Jason E. Stajich, Joseph W. Spatafora

## Abstract

Research into secondary metabolism (SM) production by fungi has resulted in the discovery of diverse, biologically active compounds with significant medicinal applications. However, the fungi rich in SM production are taxonomically restricted to Dikarya, two phyla of Kingdom Fungi, Ascomycota and Basidiomycota. Here, we explore the potential for SM production in Mucoromycota and Zoopagomycota, two phyla of nonflagellated fungi that are not members of Dikarya, by predicting and identifying core genes and gene clusters involved in SM. The majority of non-Dikarya have few genes and gene clusters involved in SM production except for the amphibian gut symbionts in the genus *Basidiobolus. Basidiobolus* genomes exhibit an enrichment of SM genes involved in siderophore, surfactin-like, and terpene cyclase production, all these with evidence of constitutive gene expression. Gene expression and chemical assays confirm that *Basidiobolus* has significant siderophore activity. The expansion of SMs in *Basidiobolus* are partially due to horizontal gene transfer from bacteria, likely as a consequence of its ecology as an amphibian gut endosymbiont.

## Introduction

Fungi produce a wealth of biologically active small molecules – secondary or specialized metabolites – that function in interactions with other organisms, environmental sensing, growth and development, and numerous other processes (Rokas et al. 2020). Several of these compounds have led to the successful development of pharmaceuticals (e.g., antibiotics, immunosuppressants, statins, etc.) that have had dramatic and positive impacts on human health. Understanding the evolution of fungal secondary metabolites and linking them with their ecological and physiological functions in nature can inform searches for compounds with applications in human society.

Secondary metabolism (SM) is imprecisely defined but can be characterized generally as the production of bioactive compounds that are not part of primary metabolism and that are not required for growth and survival in the laboratory (Keller et al 2005, Brakhage 2013, Rokas et al. 2020). In fungi, the genes responsible for the synthesis of secondary metabolites are frequently co-located in biosynthetic gene clusters (Smith et al. 1990, Brakhage 2013), which contain the genes that control regulation of expression, biosynthesis, tailoring, and transport of these compounds out of the cell (Smith et al. 1990, Keller et al 2005, Osbourn 2010). In the kingdom Fungi, the diversity of products synthesized via SM is substantial and primarily includes alkaloids, peptides, polyketides, and terpenes (Collemare et al. 2008, Helaly et al. 2018). Each of these groups of compounds are synthesized by core genes that are characteristic of the pathways and include, but are not limited to, dimethylallyl tryptophan synthases (DMAT), non-ribosomal peptide synthetases (NRPS), polyketide synthetases (PKS), and terpene cyclases (TC). These bioactive compounds fulfill various roles that are hypothesized to increase the fitness of the fungus by promoting better recognition and adaptation to environmental cues.

Biosynthesis of secondary metabolites is heterogeneous across the fungal tree of life, but the vast majority of discovered and predicted secondary metabolites are reported within the fungal phyla Ascomycota and Basidiomycota of the subkingdom Dikarya. Filamentous ascomycetes are the major producers of secondary metabolites (e.g., penicillin, cyclosporin, etc.), with the majority of genes and gene clusters involved in fungal SM discovered in the subphylum Pezizomycotina (Collemare et al. 2008, Helaly et al. 2018). Although less than Ascomycota, Basidiomycota is also a prominent producer of SM, including some of the better-known hallucinogens (e.g., psilocybin of *Psilocybe*; Reynolds et al. 2018) and compounds toxic to humans (e.g., amanitin of *Amanita*; Luo et al. 2010).

For reasons that are unclear, the remainder of kingdom Fungi is characterized by a paucity of secondary metabolites (Voight et al. 2016). This includes the zoosporic fungi and relatives classified in Blastocladiomycota, Chytridiomycota and Rozellomycota, and the nonflagellated, zygomycete fungi of Mucoromycota and Zoopagomycota. This pattern of SM diversity supports the hypothesis that diversification of secondary metabolism is a characteristic of Ascomycota and Basidiomycota (subkingdom Dikarya), which share a more recent common ancestor relative to the other phyla. Recent genome sampling efforts have focused on increased sequencing of non-Dikarya species (Nagy et al. 2014, Kohler et al. 2015, Spatafora et al. 2016, Quandt et al. 2017, Ahrendt et al. 2018). These efforts have provided a better understanding of the relationships of the phyla of kingdom Fungi (e.g., Spatafora et al. 2016), and processes and patterns that shaped the evolution of morphologies (e.g., Nagy et al. 2014) and ecologies (e.g., Chang et al. 2019, Quandt et al. 2017) within the kingdom. The availability of a diversity of these genomes provides an opportunity to characterize and focus on the secondary metabolism composition of non-Dikarya taxa, which have remained relatively unexplored.

While the majority of non-Dikarya taxa have low SM diversity, genomic sequencing of the genus *Basidiobolus* (Phylum Zoopagomycota) revealed that it possesses an unusually large composition of SM gene clusters. Species of *Basidiobolus* have complex life cycles, which produce multiple spore types that occur in multiple environmental niches. These species are symbionts found in the digestive tracts of reptiles and amphibians, but their function and impact on the host remains unknown. The fungus is dispersed with the feces where it sporulates producing both forcibly discharged asexual spores (blastoconidia) and passively dispersed asexual spores (capilloconidia) that adhere to exoskeletons of small insects. These insects are consumed by insectivorous amphibians, completing the life cycle. *Basidiobolus* also reproduces sexually through the production of zygospores (meiospores) either by selfing (homothallic) or outcrossing (heterothallic) according to species. The fungus is also isolated from leaf litter and can be maintained in pure culture, findings that are consistent with a saprobic (decomposition of organic matter) phase to the life cycle. *Basidiobolus* must have adapted for survival in numerous environmental niches including the amphibian digestive system, amphibian feces, insect phoresis, and on decaying plant matter or leaf litter.

In this study we demonstrate that the genomes of *Basidiobolus* contain a larger number of genes related to SM than predicted by phylogeny and that in several cases the evolution of many of these SM genes is inconsistent with vertical evolution. Our objectives were to: i) characterize the diversity of SM in *Basidiobolus*, ii) identify the phylogenetic sources of this diversity, and iii) determine which classes of SM gene clusters are functional and may predict the secondary metabolites produced by species of *Basidiobolus*. Finally, we propose a model in which the amphibian gastrointestinal system is an environment that promotes noncanonical evolution of its fungal inhabitants.

## Results

### Secondary metabolite gene cluster prediction

A total of 38 secondary metabolism (SM) gene clusters and 44 SM core genes were predicted in the *B. meristosporus* CBS 931.73 genome, 40 SM gene clusters and 44 SM core genes for *B. meristosporus* B9252, and 23 SM gene clusters and 23 SM core genes for *B. heterosporus* B8920 (Table 1; Supplementary Table 2). Seventy-eight percent of the SM gene models predicted were found to be shared across the *Basidiobolus* isolates. Ten SM core genes were found to be unique to *B. meristosporus* CBS 931.73, 10 SM genes unique to *B. meristosporus* B9252, and 5 SM genes unique to *B. heterosporus* B8920 (Additional file 1).

**Table 1.** Predicted secondary metabolite (SM) core genes for *Basidiobolus* isolates used in this study. NRPS: Non-ribosomal peptide synthetases, PKS: Polyketide synthases, TC: Terpene cyclases.

| Isolate | Total SM | NRPS/NRPS-like | PKS/PKS-Like | NRPS-PKS hybrids | TC |
| --- | --- | --- | --- | --- | --- |
| *B. meristosporus* CBS 931.73 | 44 | 30 | 4 | 0 | 10 |
| *B. meristosporus* B9252 | 43 | 29 | 7 | 1 | 6 |
| *B. heterosporus* B8920 | 23 | 18 | 1 | 0 | 4 |

A total of 721 SM gene clusters were predicted for the 66 additional Mucoromycota and Zoopagomycota genomes including 74 non-ribosomal peptide synthetases (NRPS), 167 NRPS-like, 97 polyketide synthases (PKS), 91 PKS-like, 292 terpene cyclase (TC) gene models across both phyla (Figure 1, Supplementary Table 2). For Zoopagomycota (including *Basidiobolus*), 284 SM gene models were predicted, including one NRPS-PKS hybrid, 71 NRPS, 56 NRPS-Like, 54 PKS, 46 PKS-Like, and 56 TC gene models. In Mucoromycota, 563 total SM gene models were predicted, including 45 NRPS, 142 NRPS-Like, 49 PKS, 51 PKS-Like, and 256 TC gene models. The three isolates with the most numerous predicted SM gene clusters, not including *Basidiobolus* genomes, were *Dimargaris cristalligena* RSA 468 with 33 predicted SM proteins (21 NRPS, 7 NRPS-Like, 2 PKS-Like, 3 TC), *Linderina pennispora* ATCC 12442 V 1.0 with 23 SM predicted (1 NRPS, 15 PKS, 3 PKS-like, 4 TC), and *Martensiomyces pterosporus* CBS 209.56 v1.0 with 23 SM proteins predicted (1 NRPS, 5 PKS, 14 PKS-Like, 3 TC). No DMAT gene models were predicted for any member of Mucoromycota or Zoopagomycota.

**Figure 1.**
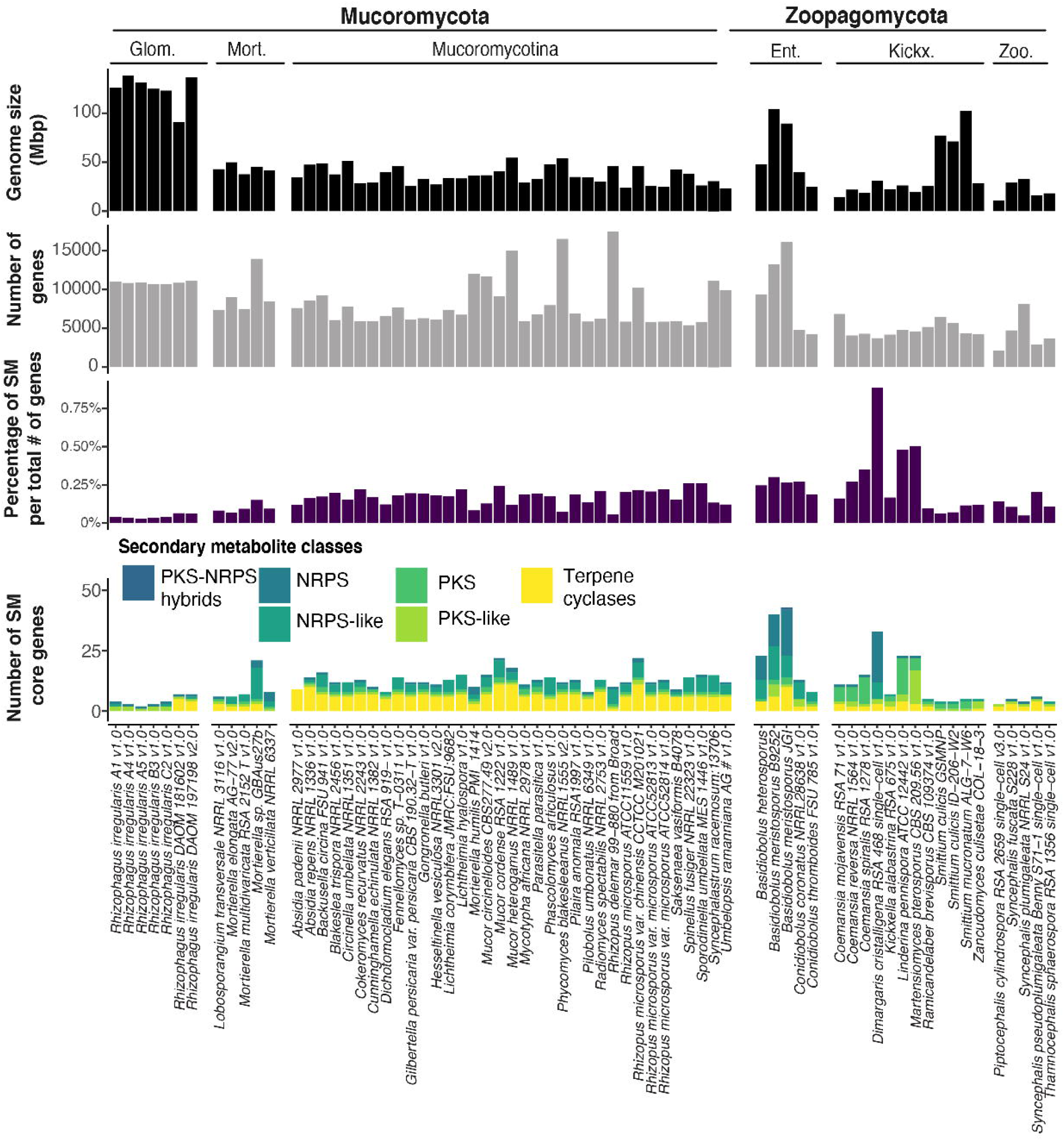
Genome size, number of genes, proportion of secondary metabolism (SM) gene clusters and number of predicted SM gene clusters for 66 Zoopagomycota and Mucoromycota sequenced genomes. Color code represents the category of SM predicted per isolate. NRPS: Non-ribosomal peptide synthetases. PKS: Polyketide synthases. Glom.: Glomeromycotina. Mort.: Morteriellomycotina. Ent.: Entomophtoromycotina. Kickx.: Kickxellomycotina. Zoo.: Zoopagomycotina

### Expression of core SM genes in Basidiobolus

A total of 83.45% of the RNA sequenced reads were mapped uniquely to the reference genome of *B. meristosporus* CBS 931.73, while 12.08% of the reads were mapped in more than one location. Only 4.47% of the RNA sequenced reads did not map to the reference genome. The majority of predicted SM for *B. meristosporus* CBS 931.73 were expressed at the same or higher levels than constitutive housekeeping genes, such as Beta-tubulin, Elongation Factor 1, Actin, and Ubiquitin (Figure 2). The highest expressed SM core genes per SM group were: NRPS – gene model 387529 (Cluster 5) with 74.03 transcripts per million (TPM) mapped; NRPS-like – gene model 221915 (Cluster 20) with 45.58 TPM mapped; PKS – gene model 290138 (Cluster 37) with 687.55 TPM mapped; PKS-like – gene model 207695 (Cluster 38) with 26.15 TPM mapped, and Terpene cyclase – gene model 301341 (Cluster 13) with 78.26 TPM mapped.

**Figure 2.**
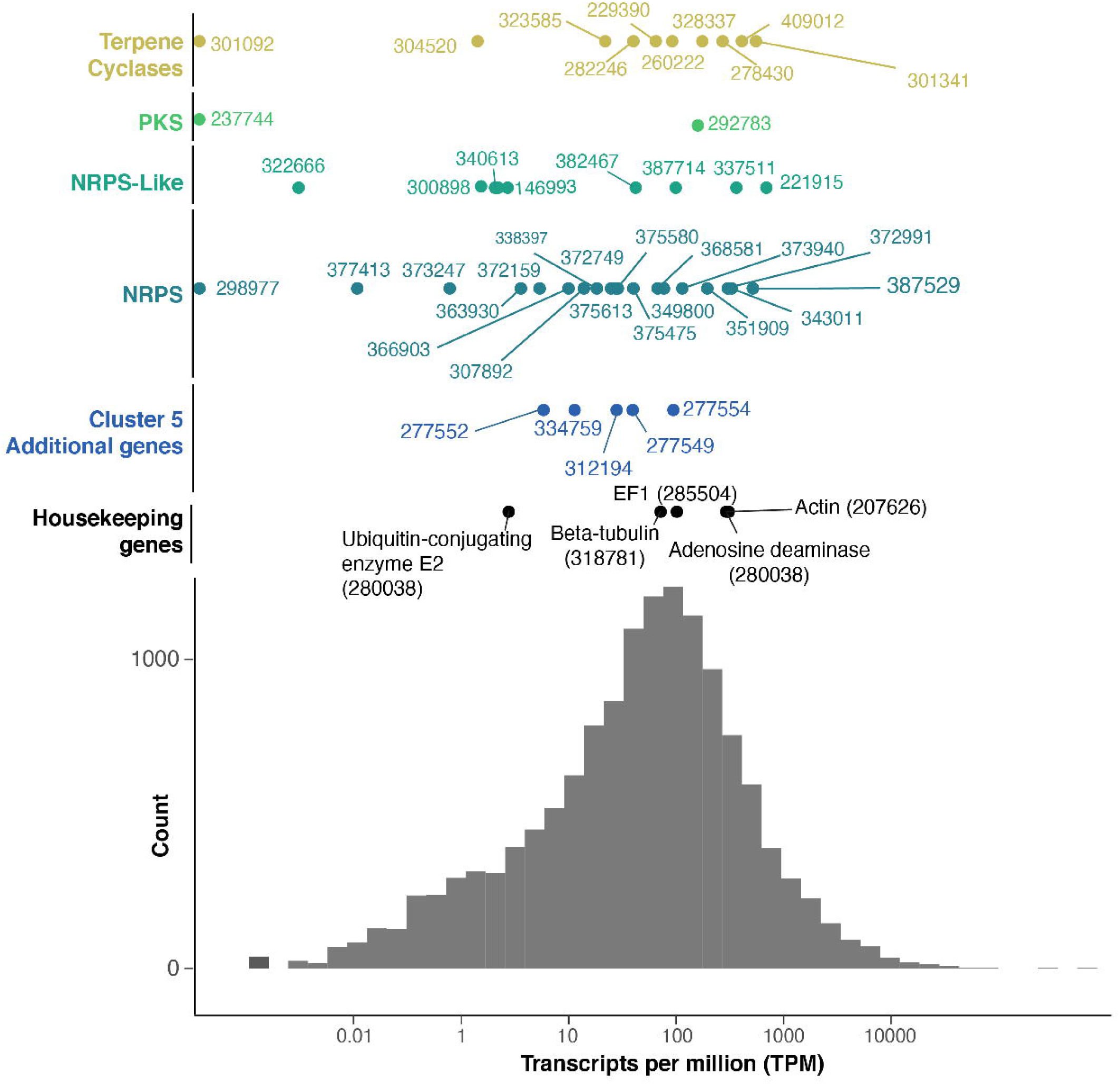
Distribution of number of RNAseq counts in transcripts per million (TPM) per genie feature from *B. meristosporus* CBS 931.73. Colors represent SM and genes of interest. X-axis represents the TPM count. Y-axis represents distribution of TPM. The scale is in log(TPM), and the values are in TPM absolute values. Histogram represents the distribution of mapped reads in TPM for all predicted gene models with non-zero TPM values across the *B. meristosporus* CBS 931.73 genome.

### Phylogenetic analysis of NRPS/NRPS-Like A-domains

The phylogenetic reconstruction of A-domains was performed with 951 A-domains from a combined dataset including the A-domain dataset from Bushley and Turgeon (2010) and the predictions of NRPS/NRPS-like A-domains from Mucoromycota and Zoopagomycota genome sequences (Additional file 2). A total of 395 NRPS A-domains were predicted for *Basidiobolus*, Mucoromycota and Zoopagomycota genome sequences (Additional File 3). The phylogenetic analyses recovered the nine major families reported by Bushley and Turgeon (2010) with the addition of the two new clades, surfactin-like and ChNSP 12-11-like, reported here. The total number of Mucoromycota/Zoopagomycota A-domains and their distribution across these clades are as follows: 74 to the AAR clade; 115 to the major bacterial clade (MBC), including 104 A-domains with the surfactin-like clade with and an additional 11 A-domains scattered elsewhere in the MBC; the CYCLO clade with eight A-domains; 65 A-domains to the ChNSP 12-11-like clade; 25 A-domains to the ChNSP 12-11 clade; 11 A-domains to the PKS/NRPS clade; 16 A-domains to the SID clade; 76 A-domains to the SIDE clade; and 5 A-domains to the EAS clade (Figure 3). No A-domains were clustered into the ACV clade or the ChNSP 10 clade, and the remaining A-domains grouped within the outgroup clade (Figures 3 and 4, Supplementary Figure 1). Similar phylogenetic origins for multiple A-domains were found in a single NRPS core gene (Figure 4).

**Figure 3.**
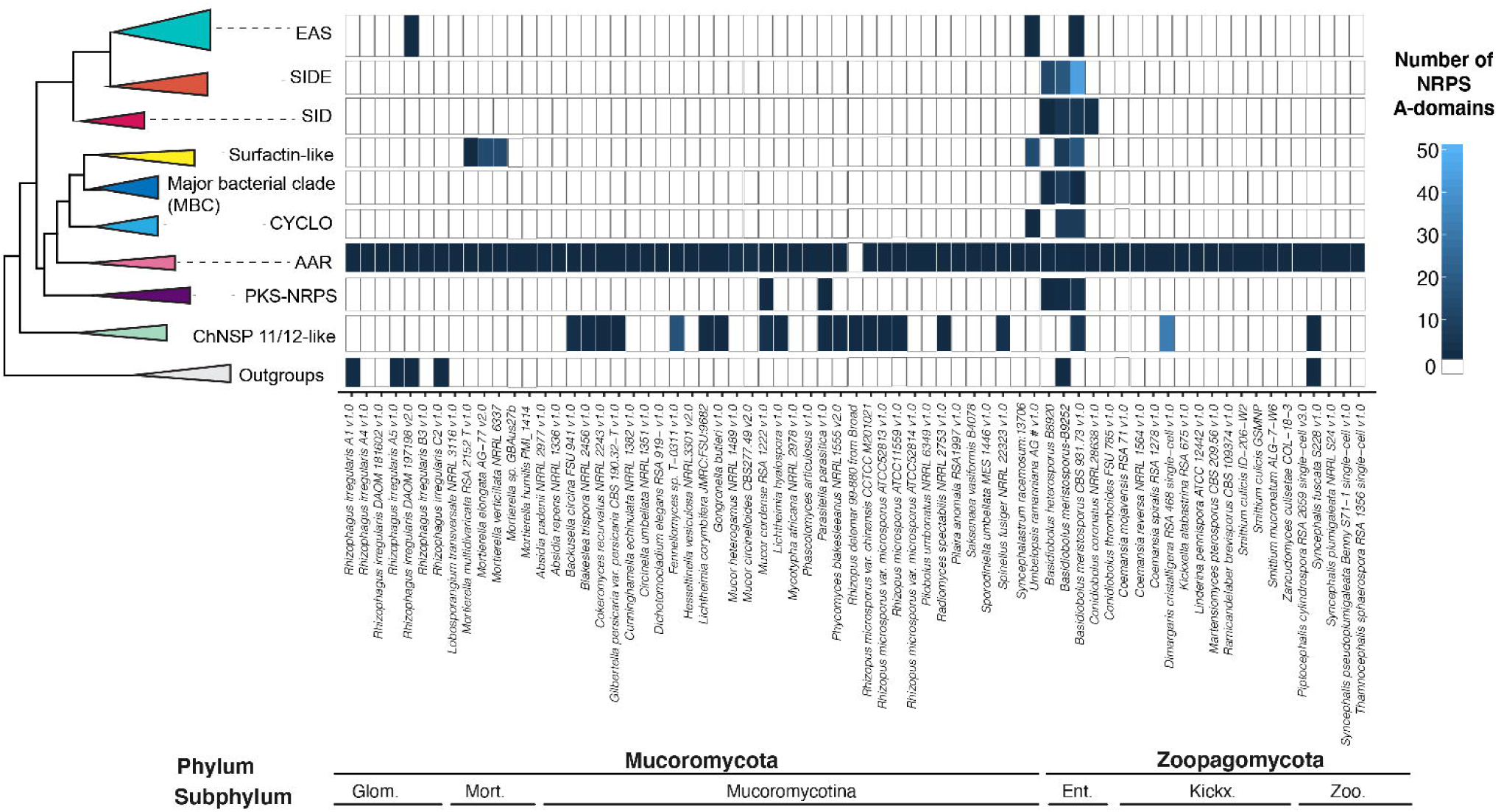
Phylogenetic sources and abundance of A-domains from NRPS predicted gene models for Zoopagomycota and Mucoromycota species. The maximum likelihood phylogenetic tree (left) represent a simplification of the reconstructed tree which includes the clades that include more than one A-domain from Mucoromycota/Zoopagomycota genomes. The heatmap (right) represents the abundances A-domains predicted for each domain clustered within each clade. *Basidiobolus* species are enriched in NRPS from the major bacterial clade (MBC), cyclosporin (CYCLO), surfactin-like, and siderophore (SIDE, SID) clades.

**Figure 4.**
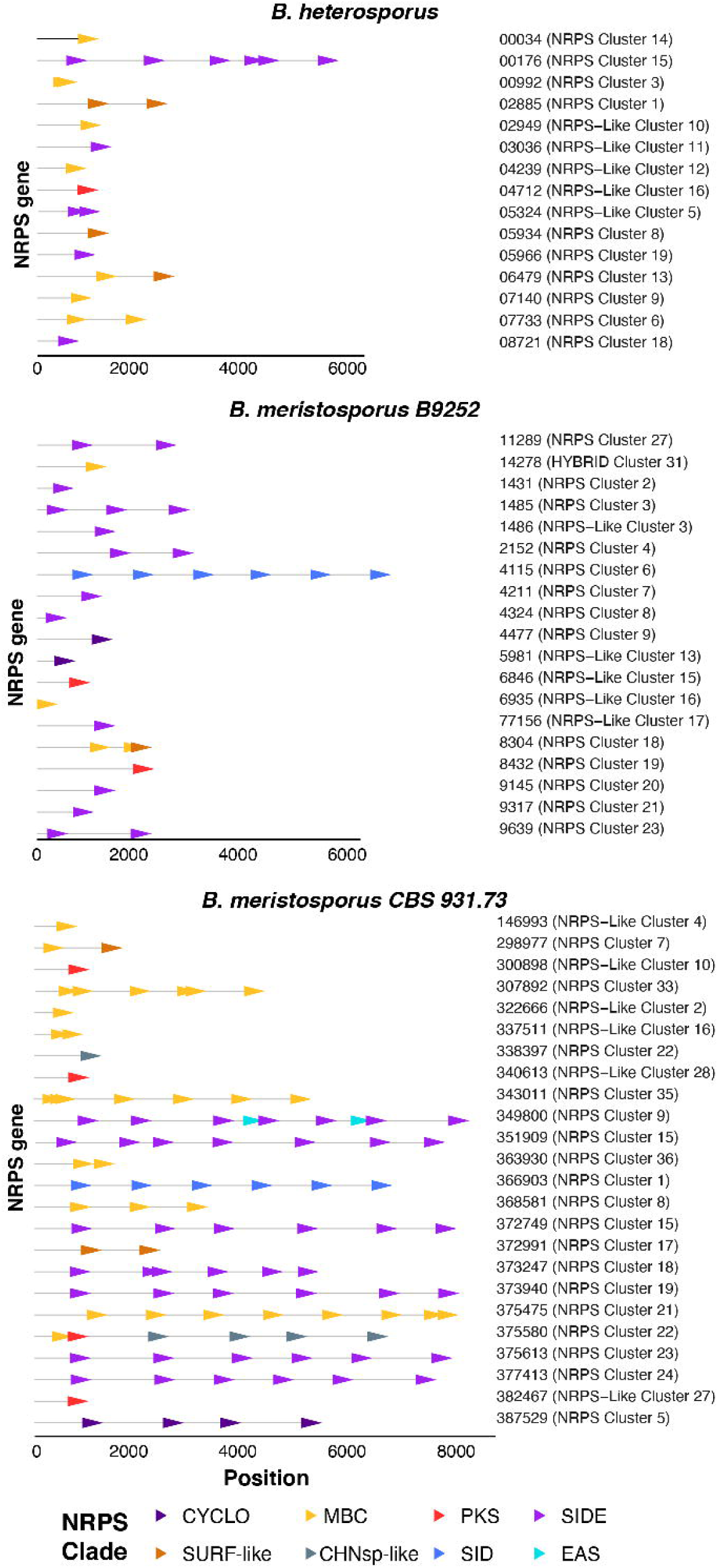
Graphical representation of the A-domains of each NRPS core gene predicted for *Basidiobolus* genomes. Horizontal grey lines represent the length of the predicted NRPS core gene. Arrows represent A-domains and are located in the position within the gene model. Colors represent the phylogenetic origin. Numbers represent the name of each gene model and predicted SM cluster for each genome.

Of fungi classified in Mucoromycota or Zoopagomycota, *Basidiobolus* genomes contained the most A-domains clustered in the EAS, SIDE, and CYCLO clades. The A-domains clustered within EAS represent two domains from *B. meristosporus* CBS 931.73, and one A-domain of *Rhizophagus irregularis* DAOM 197198 v2.0. The A-domains clustered within SIDE represent 76 domains, 44 from *B. meristosporus* CBS 931.73, 19 from *B. meristosporus* B9252, and 13 from *B. heterosporus*. The A-domains clustered within the CYCLO clade represent six domains, four from *B. meristosporus* CBS 931.73 and two from *B. meristosporus* B9252. The NRPS A-domains grouped in the CYCLO cluster correspond to cluster 5 NRPS from *B. meristosporus* CBS 931.73 (Protein ID 387529) and cluster 9 of *B. meristosporus* B9252 (Protein ID N161_4477). Gene model 387529 was predicted as a tetra-modular protein, with two N-methyltransferase domains and a thioesterase domain. In addition, AntiSMASH analyses predict a six gene cluster (cluster 5) that includes gene model 387529, a zinc finger transcription factor (312194), carrier protein (277549), transporter (277552), peptidase (277554), and a SNARE associated protein (334759) (Supplementary Figure 2). All of these gene models have evidence of gene expression (Figure 2).

The *Basidiobolus* NRPS A-domains clustered in the surfactin-like clade included 32 A-domains from eight gene models from *B. meristosporus* CBS 931.73. These included gene model 375475 (Cluster 19) with eight A-domains, gene models 307892 (Cluster 33) and 343011 (Cluster 35) with seven A-domains each; gene model 368581 (Cluster 8) with three A-domains; gene models 337511 (Cluster 16), 372991 (Cluster 17), and 298977 (Cluster 7) each with two A-domains; and gene model 322666 (Cluster 2) with one A-domain. *B. meristosporus* B9252 contained two surfactin-like A-domains from one gene model (N161_8304; Cluster 18). *B. heterosporus* possessed 13 A-domains from seven gene models, including N168_07733 (Cluster 6), N168_06479 (Cluster 13), N168_02885 (Cluster 1) and N168_00034 (Cluster 14) with two A-domains each, and gene models N168_05934 (Cluster 8), N168_04239 (Cluster 12) and N168_07140 (Cluster 9) with one A-domain each.

### Evolutionary relationships of predicted PKSs

A total of 421 KS domains were included in the phylogenetic reconstruction (Additional File 4). Two KS domains were predicted from Chytridiomycota genomes, 21 from Neocallimastigomycota, 46 from Basidiomycota, 78 from Ascomycota, 54 from Mucoromycota, and 76 from Zoopagomycota for a total of 310 predicted fungal KS domains from genome sequences (Additional File 4). The additional KS domains were obtained from Kroken et al. (2003). For Mucoromycota and Zygomycota, the genomes with the highest number of KS domains were *Linderina pennispora* ATCC 12442 v1.0 (20 KS domains), *Coemansia spiralis* RSA 1278 v1.0 (11 KS domains), *Martensiomyces pterosporus* CBS 209.56 v1.0 (10 KS domains), *Coemansia reversa* NRRL 1564 v1.0 (8 KS domains), *Coemansia mojavensis* RSA 71 v1.0 (7 KS domains), and the *Basidiobolus* genomes: *B. meristosporus* CBS 931.73 (5 KS domains), *B. meristosporus* B9252 (4 KS domains), and *B. heterosporus* (3 KS domains).

Five major clades of PKS proteins were reported by Kroken et al. (2013) and recovered by this study, including the fungal reducing (R) PKS, animal fatty acid synthases (FAS), fungal non-reducing (NR) PKS, bacterial PKS, and fungal FAS (Supplementary Figures 3 and 4). The Fungal reducing PKS1 clade included domains from *B. meristosporus* CBS 931.73 (2 KS domains) and *B. meristosporus* B9252 (2 KS domains), as well as a new sub-clade called “reducing PKS clade V”. This clade comprised KS domains from Basidiomycota, Neocallimastigomycota and one Chytridiomycota representative. No KS domains of Mucoromycota or Zoopagomycota genomes were clustered within the animal FAS clade. The Fungal NR PKS clade contained a new clade, “non-reducing PKS proteins clade IV”, comprising KS domains from Basidiomycota, Neocallimastigomycota, and one KS domain of *B. meristosporus* B9252. The Bacterial PKS clustered KS domains from *B. meristosporus* CBS 931.73 (1 KS domain) and *B. meristosporus* B9252 (1 KS domain) as sole representative of Mucoromycota and Zoopagomycota. Finally, the fungal FAS clade comprised all the remaining KS domains of Mucoromycota and Zoopagomycota genomes (Supplementary Figures 3 and 4).

To better understand the predicted PKSs of Mucoromycota and Zoopagomycota that clustered in the fungal FAS clade, the patterns of domains that comprise fungal PKS protein sequences were analyzed. Predicted PKS of Ascomycota contained AT, KS and PP domains in all sequences (Supplementary Figure 5). Predicted PKS from Basidiomycota contained AT and KS domains in all sequences, while KR and PP domains were present on 36% of the sequences. All PKS that contain a KS domain in the FAS clade are missing the PP domain. For Chytridiomycota, the predicted PKS with the KS domain clustered in the Fungal (R) clade contained AT, KS and DH domains, but no PP domain. Chytridiomycota PKS with KS domains associated to FAS clade only possessed AT and KS domains. For Neocallimastigomycota, 100% of PKS clustered in fungal R and NR clades contained AT and KS domains, however, only 5 PKS contained the PP domain. All Neocallimastigomycota PKS that clustered within the FAS clade contained only AT and KS domains. For Mucoromycota and Zoopagomycota, 100% of PKS contained either AT and/or KS domain, but no other domains (Supplementary Figure 6).

### Evolutionary relationships of predicted terpene cyclase gene models

A total of 1,108 terpene cyclase or terpene cyclase-like gene models were identified and used for phylogenetic analyses (Additional file 5). These include 256 identified in the Mucoromycota species genomes, 56 in the Zoopagomycota genomes, and 401 ortholog proteins from 58 fungal genomes of Basidiomycota and Ascomycota. An additional 395 TC candidates were identified in bacterial genomes from RefSeq via BLASTP. The phylogenetic reconstruction of TC resulted in two main clades: an outgroup clade that comprised predicted TC annotated as tRNA threonylcarbamoyladenosine dehydratase and phytoene synthases, and a main ingroup TC core clade (Figure 5, Supplementary Figure 7). The tRNA threonylcarbamoyladenosine dehydratase subclade contained mostly bacterial sequences plus one TC gene model of *B. meristosporus* B9252 and *B. meristosporus* CBS 931.73 each. The phytoene synthases clade comprised two subclades, one clade that grouped most bacterial gene models, and a second clade clustering bacterial and Mucoromycota predicted TC gene models, but no *Basidiobolus* TC core gene models were found in this clade.

**Figure 5.**
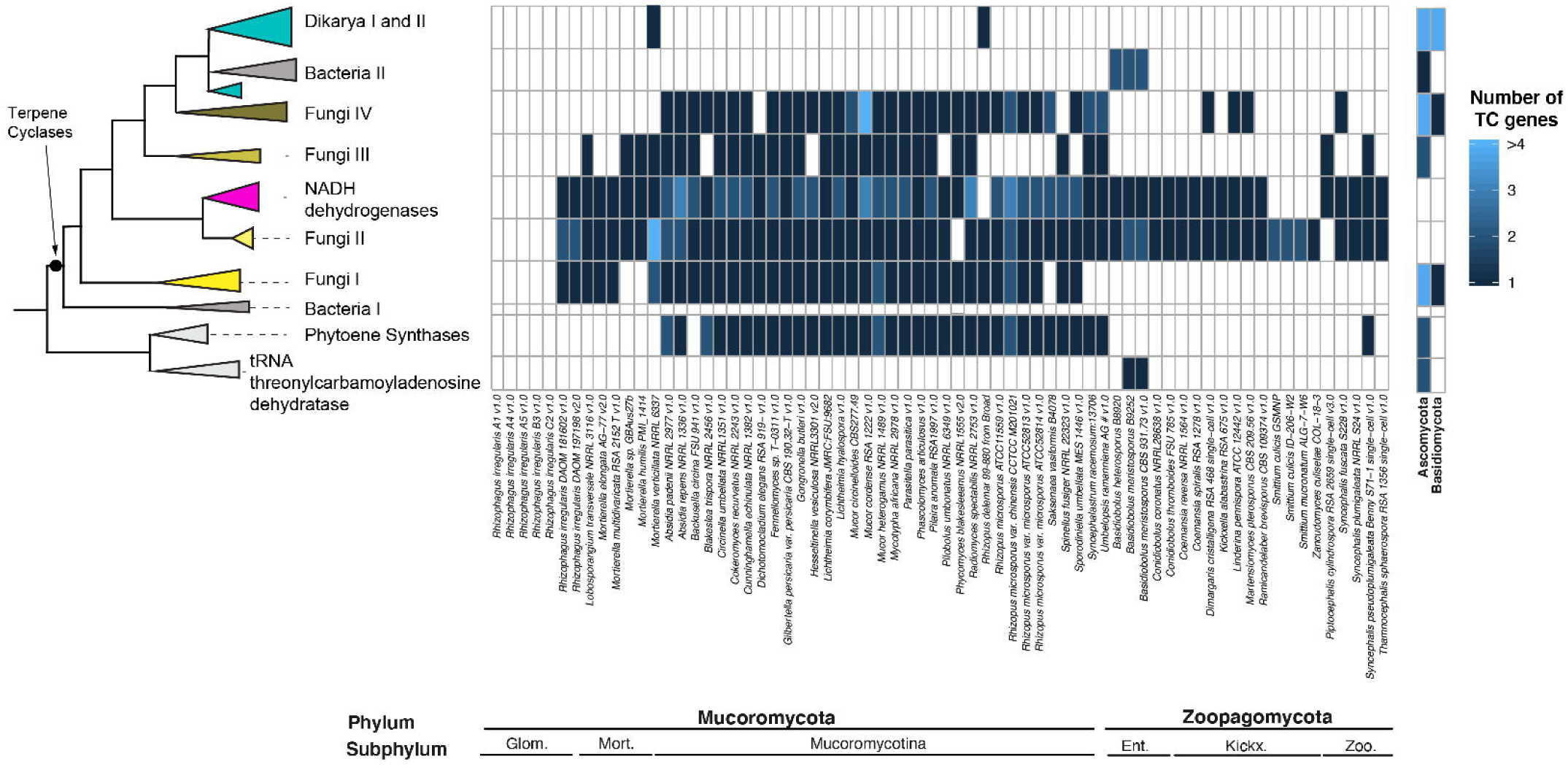
Phylogenetic sources and abundance ofterpene cyclase (TC) core genes predicted for Zoopagomycota and Mucoromycota species. The maximum likelihood phylogenetic tree is a simplification of the maximum likelihood tree reconstructed using the TC core genes. The heatmap (right) represents the abundances of predicted TC genes within each reconstructed clade. Glom.: Glomeromycotina. Mort.: Morteriellomycotina. Ent.: Entomophtoromycotina. Kickx.: Kickxellomycotina. Zoo.: Zoopagomycotina

The TC core clade (Figure 5 and Supplementary Figure 7) comprised a bacterial exclusive clade (Bacteria I), followed by four highly supported sub-clades containing TC core genes from Mucoromycota, Zoopagomycota, and Dikarya, and are referred to here as Fungi TC clades I – IV. The majority of TC genes from Mucoromycota and Zoopagomycota clustered within clade Fungi TC II, which included no other fungal or bacterial TC. These genes were annotated as Squalene synthetases by the *Mycocosm* genome portal.

The Fungi TC clades I, III and IV included mostly Mucoromycota TCs. The Fungi TC I clade contained TC genes from Mucoromycota isolates, as well as TC genes from the ascomycete isolates *Fusarium verticillioides* 7600 and *Hypoxylon sp*. EC38 v1.0, and the basidiomycete *Suillus luteus* UH-Slu-Lm8-n1 v2.0. These gene models were annotated as associated with the Ubiquitin C-terminal hydrolase UCHL1 according to the *MycoCosm* portal. Fungi TC III clade also comprised mostly Mucoromycota isolates, with additional TCs from Zoopagomycota isolates *Piptocephalis cylindrospora* RSA 2659 single−cell v3.0 and *Syncephalis pseudoplumigaleata* Benny S71−1 single−cell v1.0. Annotations of genes in Fungi TC III clade in *Mycocosm* indicated that these proteins are part of the isoprenoid/propenyl synthetases, responsible for synthesis of isoprenoids. Isoprenoids play a role on synthesis of various compounds such as cholesterol, ergosterol, dolichol, ubiquinone or coenzyme Q (Finn et al. 2009). Finally, Fungi TC IV clade followed a similar Mucoromycota-enriched pattern with the exceptions of *Dimargaris cristalligena* RSA 468 single−cell v1.0, *Linderina pennispora* ATCC 12442 v1.0, *M. pterosporus* CBS 209.56 v1.0, and *Syncephalis fuscata* S228 v1.0. Fungi TC IV clade also included TC genes from Ascomycete sequenced genomes. The majority of these genes were annotated as containing a terpene synthase family, metal binding domain, and a polyprenyl synthetase domain in the *MycoCosm* genome portal.

Within the fungal clades, the NADH dehydrogrenase (ubiquinone) complex clade represented a highly supported clade clustering bacterial, Mucoromycota and Zoopagomycota TCs. Annotations associated with gene models found in this clade indicated that this clade comprised TC associated to NADH dehydrogrenase (ubiquinone) complex.

The terminal clades of TC clusters include the Bacteria II + *Basidiobolus* clade that included bacterial TC and six TC SM core genes predicted from *Basidiobolus* (two from each genome), and the clade comprising TC core genes solely from Dikarya and one gene from *Mortierella verticillata NRRL 6337* and *Rhizopus microsporus ATCC11559 v1*.*0* (Figures 5 and Supplementary Figure 7).

### *Signatures for HGT in* Basidiobolus *genomes*

The identity search for genes with evidence for HGT identified 934 genes in *B. meristosporus* CBS 931.73, 620 genes of *B. meristosporus* B9252, and 382 genes of and *B. heterosporus* B8920 with zero BLASTP hits to fungal proteins. These genes were used for the coverage assay to identify gene model coverage deviation from the harboring scaffold median coverage (Sup. Fig. 7). This assay resulted in 810 genes of *B. meristosporus* CBS 931.73, 503 genes of *B. meristosporus* B9252, and 301 genes of and *B. heterosporus* B8920 with z-scores under 2 standard deviations from the harboring scaffold median coverage. These genes were considered candidate genes with evidence for HGT, and represented 5%, 4% and 3% of the gene content of *B. meristosporus* CBS 931.73, *B. meristosporus* B9252, and *B. heterosporus* B8920, respectively. The HGT candidates showed significant differences in GC content when contrasted to genes considered of fungal origin (Mean fungal GC content (expected) = 0.492, Mean HGT GC content (Observed) = 0.486, t = 3.4119, df = 891.77, p-value = 0.0006741), but no differences in codon usage or 5-mer composition were detected between HGT candidate and fungal genes (Supplementary Figures 10 and 11). Differences were also found for intron number and normalized intron length between genes HGT candidate genes and fungal genes, where the distribution of intron number shows a median of 0 introns for HGT candidates and 2 for fungal genes (Kruskal-Wallis test; *χ*^2^=272.88, df = 1, p-value < 2.2e-16; Sup. Fig. 9). The majority of HGT candidate genes (61% of the genes) have no introns, compared to 36% of genes from fungal origin with no introns (Sup. Fig. 9). The candidate HGT genes have a significantly smaller normalized intron length than the genes with a fungal origin (Kruskal-wallis test; *χ*^2^=272.88, df = 1, p-value < 2.2e-16).

A large percentage of SM core genes appeared to be the product of HGT from bacterial species into *Basidiobolus*. The SM core genes identified as candidate HGT genes is 61%, 44% and 52% for *B. meristosporus* CBS 931.73, *B. meristosporus* B9252, and *B. heterosporus* B8920, respectively (Table 2). NRPS/NRPS-like SM core genes represent the largest percentage of HGT evidence, while TC have the lowest percentage of HGT evidence (Table 2, Table 3). The identification of taxonomic sources for HGT into *Basidiobolus* indicated that most HGT comes from bacteria in the phylum Proteobacteria with 234, 160 and 95 gene models in *B. meristosporus* CBS 931.73, *B. meristosporus* B9252, and *B. heterosporus* B8920, respectively. Firmicutes (112, 63, and 39 gene models, respectively) and Actinobacteria (127, 67, and 40 gene models, respectively) (Figure 6, Supplementary Table 4) were the second and third most abundant source of HGT from bacteria into *Basidiobolus*.

**Table 2.** Number of predicted secondary metabolite core genes with evidence for HGI in *Basidiobolus* genomes. NRPS: Non-ribosomal peptide synthetases, PKS: Polyketide synthases, TC: Terpene cyclases.

| Isolate | Total SM | NRPS/NRPS-like | PKS/PKS-Like | NRPS-PKS hybrids | TC |
| --- | --- | --- | --- | --- | --- |
| *B. meristosporus* CBS 931.73 | 26/44<br>(61%) | 22/30<br>(60%) | 2/4<br>(50%) | 0/0<br>(0%) | 2/10<br>(2%) |
| *B. meristosporus* B9252 | 19/43<br>(44%) | 16/29<br>(55%) | 1/7<br>(14%) | 0/1<br>(0%) | 2/6<br>(1.4%) |
| *B. heterosporus* B8920 | 12/23<br>(52%) | 12/18<br>(66%) | 0/1<br>(0%) | 0/0<br>(0%) | 0/4<br>(0%) |

**Figure 6.**
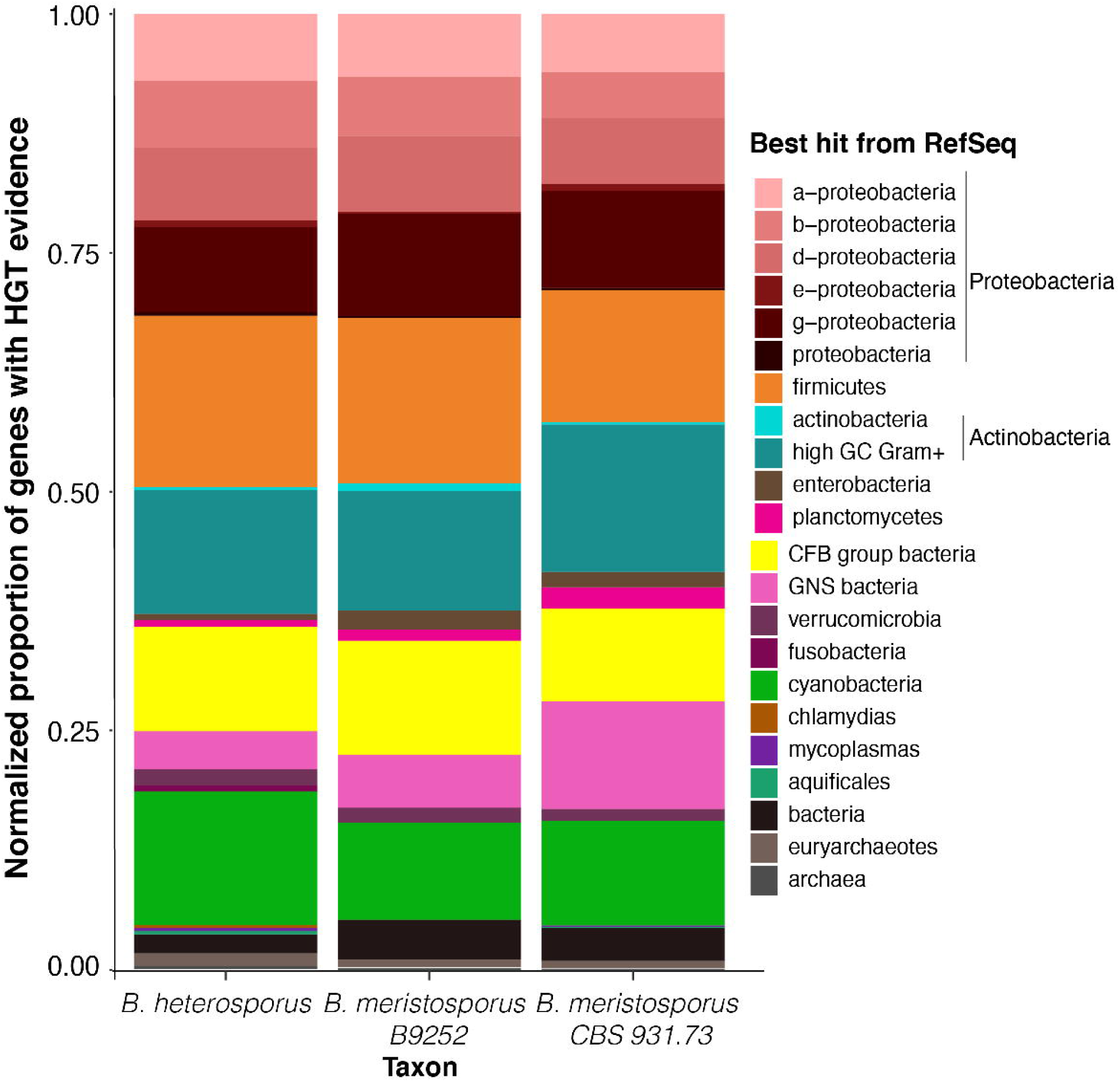
Plausible taxonomic sources of HGT genes. Bar-plot represents the proportion of diversity of HGT candidates for each *Basidiobolus* genome. Colors represent the taxonomy term for the RefSeq best hit from BLAST. Overall, between 3% to 5% of the gene models predicted for Basidiobolus species appear to be product of HGT from taxonomic groups of bacteria associated to reptilian and amphibian gut tracts (Proteobacteria, firmicutes, and CFB/bacterioidetes).

**Figure 7.**
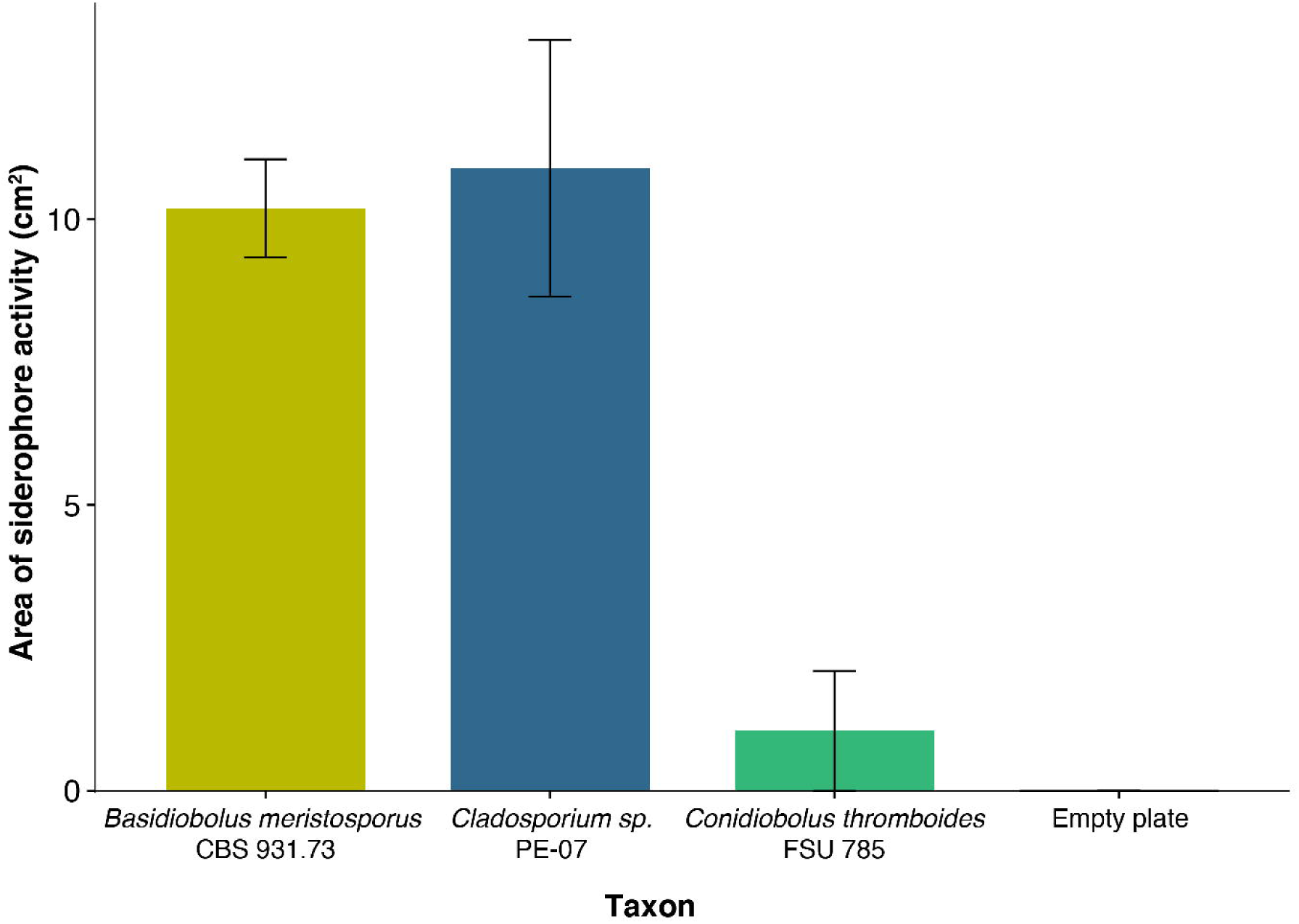
Siderophore activity of *Basidiobolus meristosporus* CBS 931.73 in a universal CAS assay using layered AY-CAS plates after 12 days. Bars represent mean siderophore activity measured per strain as the yellow area in AY-CAS plates for three replicates. Error bars represent the standard deviation for each replicate. *Cladosporium sp*. PE-07 represents the positive control. *Conidiobolus thromboides* FSU 785 represents a zygomycete with no evidence for siderophore NRPS expansion.

**Table 3.**
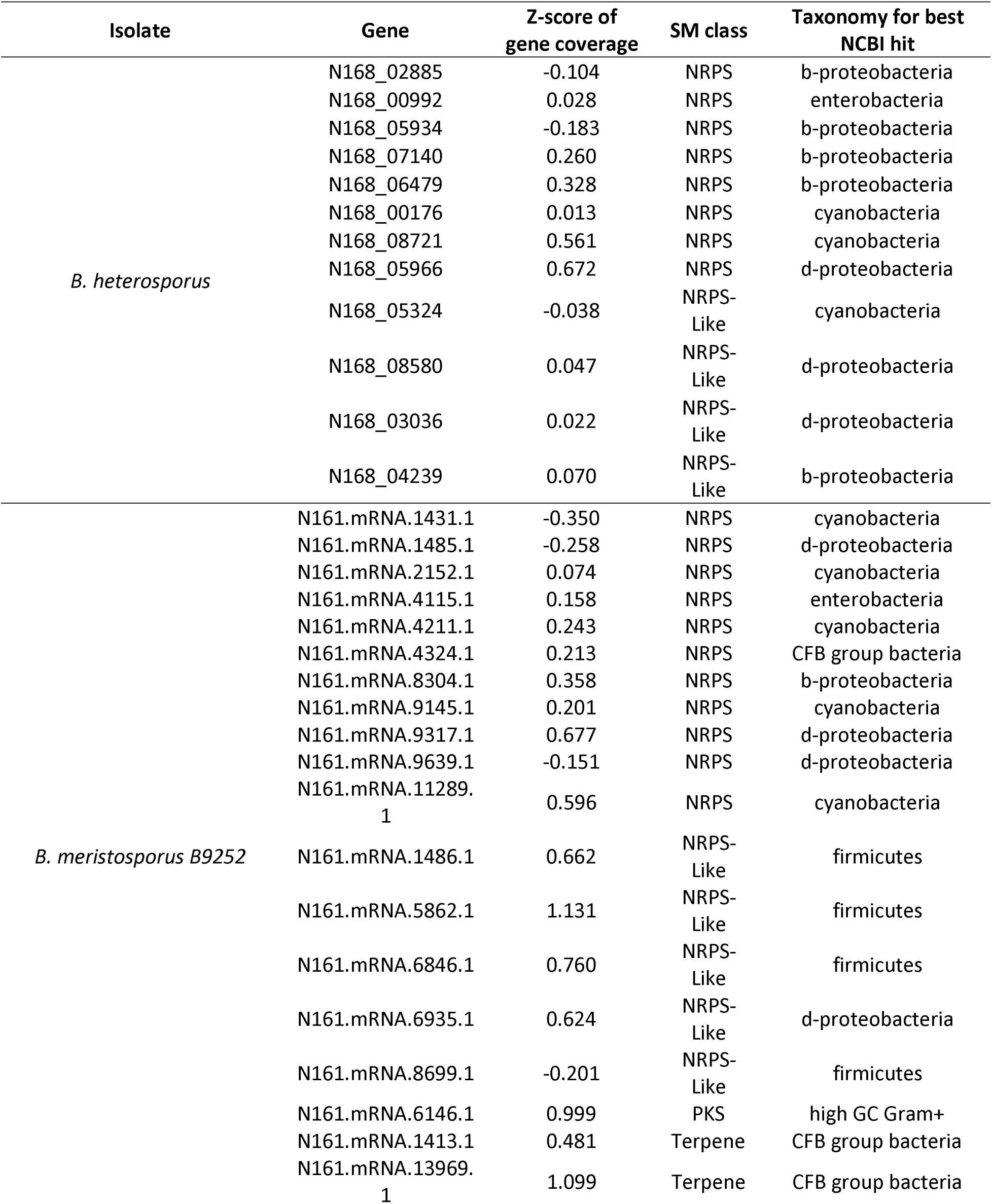
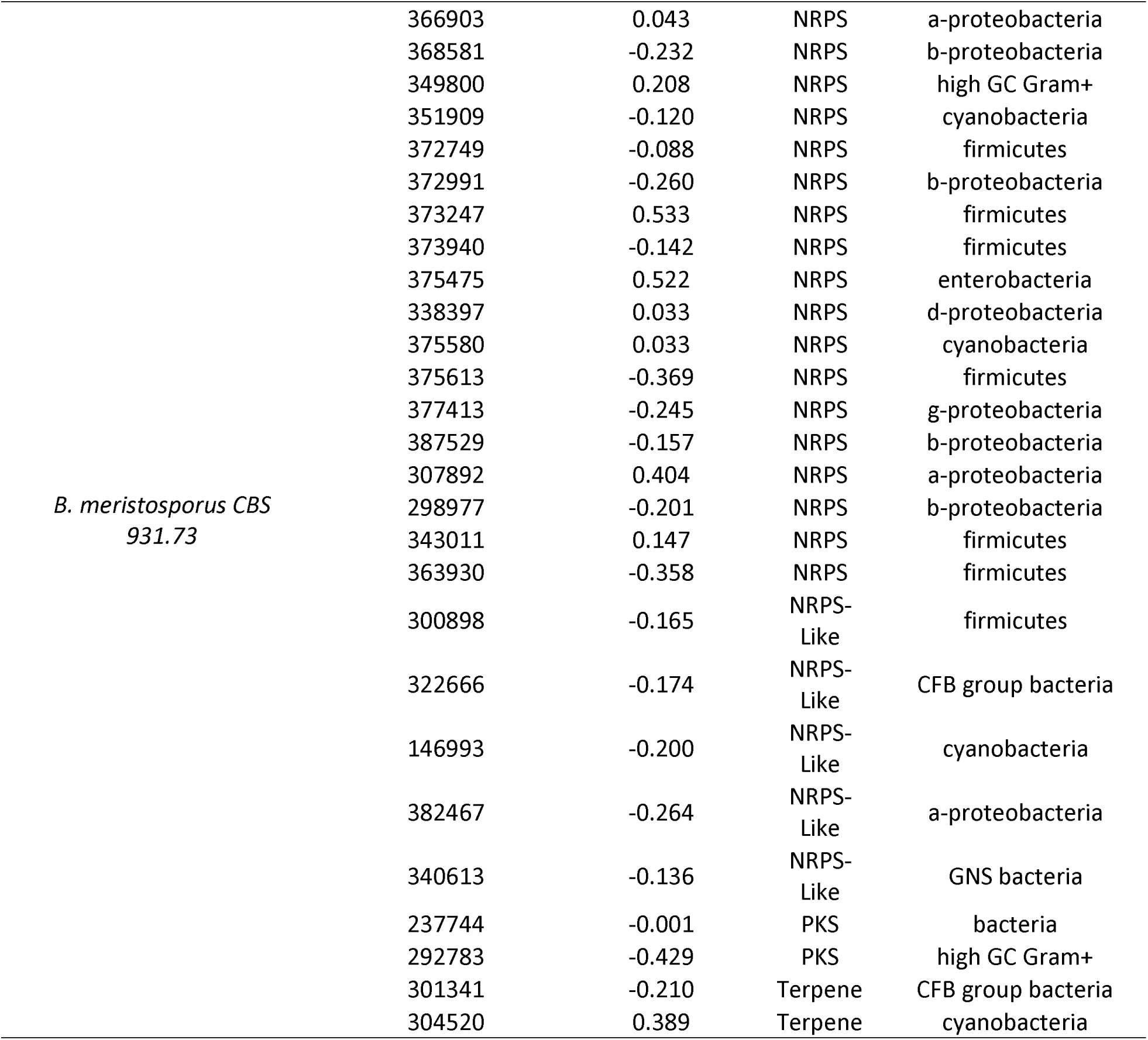
Summary of secondary metabolite core genes with HGT evidence from *Basidiobolus* isolates.

Most HGT candidates genes resulted in no gene ontology (GO) annotation (659 gene models) or no InterPro domain annotation (140 gene models, Supplementary Table 5). The ten top GO terms were oxidation-reduction process (59 gene models), protein binding (34 gene models), phosphorelay sensor kinase activity (32 gene models), N-acetyltransferase activity (28 gene models), catalytic activity (26 gene models), extracellular space (25 gene models), hydrolase activity (24 gene models), oxidoreductase activity (23 gene models), hydrolase activity (24 gene models), hydrolyzing O-glycosyl compounds(23 gene models), and NRPS (18 gene models) (Figure 6, Supplementary Table 5).

### Siderophore activity in Basidiobolus meristosporus

The siderophore activity assay resulted in visible activity for the three replicates for *B. meristosporus* and *Cladosporium sp*., and a small halo for one of the replicates of *C. thromboides*. No visible siderophore activity was detected for the other two replicates of *C. thromboides* or for the empty AY-CAS plates (Figure 7, Supplementary Figure 12). The analysis of variance for the area of siderophore activity measured showed significant differences across isolates/empty plates (ANOVA, F value = 19.62, *p* = 0.0004). Post-hoc tests showed no differences between *B. meristosporus* and *Cladosporium sp*. siderophore activity (Tukey HSD, *p* = 0.9801489) nor between *C. thromboides* and the empty AY-CAS plates (Tukey HSD, *p* = 0.9396603), but significant differences were found for all other comparisons (Tukey HSD, *p* < 0.001 for all remaining comparisons).

## Discussion

Secondary, or specialized, metabolite (SM) production is an important element of fungal metabolism. It has resulted in numerous natural products with human health implications, such as mycotoxins in our food supply, and medical applications, including antibiotics, immunosuppressants, and antitumor agents. The majority of the known genetic and chemical diversity of fungal SM has been described from the phyla Ascomycota and Basidiomycota with few SM gene clusters, and limited SM production, reported for fungi in Mucoromycota and Zoopagomycota. This observation has led to the dogma that ‘zygomycete’ species are depauperate of these chemical pathways (Voight et al. 2016). Here we report gene clusters involved in SM production in the largest survey to date of Mucoromycota and Zoopagomycota species using genomics approaches and estimate the SM potential of these fungi.

Genome sequencing of a total of 69 isolates across diverse lineages of Mucoromycota and Zoopagomycota enabled detailed identification of SM clusters. These results support the hypothesis that zygomycete fungi have a low abundance of secondary metabolism (Figure 1, Supplementary Table 2) and agree with previous reports (Voight et al. 2016). Outliers to this pattern exist, however, and are particularly true of the genus *Basidiobolus* (Zoopagomycota), which possesses a large number of SM gene clusters predicted for the NRPS, PKS and TC families. This discovery of abundant candidate genes for production of secondary metabolite in *Basidiobolus* is novel and its presence is most consistent with a signal of horizontal gene transfer from bacteria to fungi, a phenomenon we propose is facilitated by living in the amphibian gut environment.

### Distribution and Evolution of NRPS Genes Across Mucoromycota and Zoopagomycota

A deeper examination of *Basidiobolus* SM gene clusters indicates that this genus surpasses the number of expected NRPS genes for zygomycete species (Figure 1). Most of these SM genes also show evidence for transcription, indicating that the majority of these genes are expressed constitutively under laboratory conditions (Figure 2). Several of these core genes, such as the NRPS gene model 387529, appear to be expressed at a higher rate than several housekeeping genes.

A more detailed census of core genes reveals unique patterns of evolution of NRPS genes across Mucoromycota and Zoopagomycota when compared to Dikarya fungi (Figure 3, Table 1). The only NRPS A-domains found throughout the two phyla are members of the AAR clade (with the exception of *Rhizopus delemar* 99-80). These A-domains are from the genes that encode α-aminoadipate reductases (AAR), an enzyme responsible for the reduction of alpha-aminoadipic acid, which is essential for the lysine biosynthesis pathway and is present in all fungal phyla (Bushley and Turgeon 2010). In contrast, the remaining NRPS genes, and their respective A-domains, show discontinuous and patchy distributions across Mucoromycota and Zoopagomycota (Figure 3).

The most pronounced NRPS diversification in *Basidiobolus* are for genes that are predicted to encode for siderophores, iron chelating metabolites. A-domains for predicted siderophores are distributed throughout four clades including the three clades of major bacterial genes and the fungal SID and SIDE clades (Figure 3, Sup. Fig. 1). Major bacterial clades (MBC) exclusively comprise bacterial siderophore synthases, such as pyoverdine, yersiniabactin and pyochelin (Bushley and Turgeon 2010), with the exception of the surfactin-like clade. Our results show that genomes of *Mortierella* and *Basidiobolus* contain A-domains that are members of the MBC, and that they are the only fungal representatives of these clades. SID clade contains all NRPS associated with siderophore production from Ascomycota and Basidiomycota species. All *Basidiobolus* isolates contain one NRPS A-domain in this clade, as well as three A-domains from three NRPS gene models of *Conidiobolus coronatus* NRRL28638. Finally, SIDE, a clade comprising NRPS genes responsible for the production of siderophores in filamentous ascomycetes is expanded in *Basidiobolus* (Figure 3), which is also the only zygomycete with A-domains clustered in this clade. These findings are consistent with enrichment of both bacterial and fungal siderophores and are suggestive of the importance of iron metabolism in *Basidiobolus*.

The CYCLO clade contains the A-domains for core genes associated with biosynthesis of cyclic peptides, such as beauvericin and cyclosporin (Bushley and Turgeon 2010, Bushley et al 2013). Sister to all other CYCLO clade A-domains are the A-domains that comprise the NRPS core gene model 387529 from *B. meristosporus* (Supplementary Figure 2). 387529 is expressed at the highest rate of any SM gene under laboratory conditions (Figure 2). It is annotated as a tetra-modular gene model, that includes two N-methylation domains, four adenylation domains, four condensation domains, and a TE domain. When compared to *simA*, the NRPS responsible for biosynthesis of cyclosporin, structural similarities can be found, such as the presence of the N-methylation domains and the TE terminator domain. Its phylogenetic and structural similarities to *simA* suggest that 387529 results in the synthesis of a cyclic peptide with methylated amino acid residues.

The surfactin-like clade contains A-domains for bacterial core genes with similarities, but not identical, to the *Bacillus subtilis* surfactin termination module (*srfA-C* gene; Peypoux et al. 1999) including the third, fourth and fifth A-domains of the five A-domain gene model NP_930489.1 of *Photorhabdus luminescens* gene, two domains of the gene PvdD (AAX16295.1; pyoverdine synthetase) of *Pseudomonas aeruginosa*, and a single A-domain of the bimodular NRPS dbhF protein of *Bacillus subtilis* (AAD56240.1) which is involved in the biosynthesis of the siderophore bacillibactin. This surfactin-like clade contains eleven A-domains predicted from *Basidiobolus* genomes including the gene models 298977, 368581 and 372991 from *B. meristosporus* CBS 931.73. Gene model 298977 showed no evidence for gene expression, while gene models 368581 and 372991 show high rates of expression (Fig. 2). This is the first report of the prediction of a surfactin-like gene in fungi, but surfactant production was recently reported in *Mortierella alpina* (Baldeweg et al. 2019). These include malpinins, amphiphilic acetylated hexapeptides that function as natural emulsifiers during lipid secretion, and malpibaldins, hydrophobic cyclopentapeptides. This finding is consistent with the genomic data and reveals that *Mortierella*, in addition to *Basidiobolus*, possesses homologs in the surfactin-like clade that are phylogenetically different from *B. subtilis* surfactins.

Surfactins, encoded by the *srfA* gene cluster in *Bacillus subtilis*, are functionally active as surfactants, as well as toxins and antibiotics. However, the surfactin genes from *B. subtilis* (SrfA-AA, SrfA-AB and SrfA-AC) were included in our analysis and clustered in a different clade within the MBC. A-domains from single module NRPS-like protein from *B. meristosporus* CBS 931.73 (Gene model 146993) and from a NRPS-PKS hybrid A domain from of *B. meristosporus* B9252 (Gene model N161_14278) clustered with the *B. subtilis* SrfA genes. The placement of this NRPS-PKS A-domain is interesting because surfactins are lipopeptides, which contain a hydrophobic fatty acid chain, whose biosynthesis is consistent with an NRPS-PKS hybrid. The A-domains from the remaining Mucoromycota and Zoopagomycota NRPS-PKS hybrids clustered as sister to the original clade of Dikarya NRPS-PKS hybrids, supporting the hypothesis of a single origin of fungal NRPS-PKS hybrids (Bushley and Turgeon 2010).

### Mucoromycota and Zoopagomycota lack PKS diversity

Polyketide synthases (PKS) are abundant SM of Ascomycota and Basidiomycota and are involved in antibiotic production, carotenoid biosynthesis and other functional roles. In contrast, literature on PKS diversity for Mucoromycota and Zoopagomycota is limited. Our analyses update the phylogenetic reconstruction reported by Kroken et al. (2013) by adding genomic information from Chytridiomycota, Mucoromycota and Zoopagomycota (Supplementary Figures 3 and 4). Overall, we report the discovery of two new clades: A clade of non-reducing PKS proteins (clade IV) comprising Neocastimastigomycota and Basidiomycota, and a reducing PKS clade (reducing PKS clade V) consisting of Neocastimastigomycota, Chytridiomycota and Basidiomycota. The results of our PKS prediction revealed a number of potential PKS core genes in zygomycete species (Supplementary Figures 3 and 4), but the results of our phylogenetic reconstruction show that the KS domain of the predicted PKS gene models are fungal fatty acid synthases (FAS). Only *Basidiobolus meristosporus* genomes possessed KS domains associated with fungal PKS, either reducing or non-reducing.

A domain-by-domain presence/absence analysis of PKS genes models (Supplementary Figures 5 and Figure 6) shows that in addition to AT and KT domains, a third domain (either KR, DH or PP) is found in the majority of fungal PKSs (Supplementary Figure 5). Conversely, the zygomycete genes in the FAS clade only possess the AT and/or KT domains, which are domains common between FAS and PKS, including the majority of PKSs predicted for *Basidiobolus*. However, *Basidiobolus* KS domains are found in both fungal and bacterial PKS clades. Gene models 292783 and 237744 from both isolates of *B. meristosporous* cluster with fungal reducing PKS II and possess the DH domain (Supplementary Figure 4), and the expression levels of 292783 is consistent with an actively transcribed gene (Figure 2).

### Terpene cyclase-like genes are expanded in zygomycetes

Terpene cyclases (TC) are the most common SM predicted for zygomycete species (Figure 1, Supplementary Table 2). Phylogenetic reconstruction shows that zygomycete TC core genes are clearly distinct from Ascomycota and Basidiomycota TC, where at least 4 new clades of predominantly zygomycete TC are found (Figure 5, Supplementary Figure 7). Fungi II TC clade comprises TC from all zygomycete genomes analyzed with the exception of *Piptocephalis cylindrospora* RSA 2659 single−cell v3.0, *Phycomyces blakesleeanus NRRL1555* v2.0, and five genomes of *Rhizophagus irregularis*, which show no prediction of TC in their genomes. No Dikarya TC genes cluster within this clade. Fungi I, III and IV group TC genes are found almost exclusively in Mucoromycota species, as well as some Dikarya species. TC genes from Fungi I are present only in Mucoromycotina genomes. Functional annotations of TC genes from this clade indicate that these TC genes code for proteins associated with squalene and phytoene synthases and are part of the synthesis of carotenoids. Carotenoids are important compounds for the synthesis pathway of trisporic acid, the main molecule responsible for initiating sexual reproduction in zygomycetes (Burmester et al. 2007). Finally, *Basidiobolus* is the only non-Dikarya genus with TC genes clustered within the Bacteria II clade of TC. Both these SM core genes show evidence for expression comparable to housekeeping genes or other SM core genes (Figure 2). The presence of bacterial-like TC genes in *Basidiobolus* present more evidence on the plasticity of the genome of *Basidiobolus* and its ability to integrate and possibly express foreign SM associated DNA.

### Horizontal Gene Transfer of SM genes to Basidiobolus?

*Basidiobolus* is a genus with a complex biology, alternating ecologies, and multiple spore types. This complex biology is also reflected at the genomic scale. The sequenced genomes show a larger genome size than other zygomycetes, as well as a higher number of genes than any other Zoopagomycota genomes sequenced to date. Our SM prediction assay is concordant with these patterns in which *Basidiobolus* has an excess of SM gene clusters when compared to other zygomycetes. Evidence points to HGT as a main driver of SM diversity in *Basidiobolus* as supported by the phylogenetic reconstructions of NRPS, PKS and TC gene clusters with bacterial homologs. *Basidiobolus* and *Mortierella* are the only fungi with genes associated with the bacterial clades in each of these SM phylogenetic reconstructions. Moreover, these HGT candidates are integrated into the *Basidiobolus* genome assembly and do not show evidence of artifactual assembly as evidenced by discontinuous coverage (Supplementary Figure 8). The most abundant functional group of SM core genes overall are siderophores and their overall functionality is supported by both the RNA expression analyses (Figure 2) and siderophore plate assays (Figure 7).

One stage of the *Basidiobolus* life cycle the fungus lives as a gut endosymbiont where it co-occurs with bacteria and other organisms that comprise the gut microbiome. Animal gut environments can facilitate HGT between bacteria and fungi, as previously reported for the zoosporic species of Neocallimastigomycetes (Chytridiomycota), which live in the ruminant gut environment and whose genomes exhibit a 2-3.5% frequency of genes with HGT evidence (Wang et al. 2018; Murphy et al. 2019). Phylogenomic analyses of the *Basidiobolus* NRPS A-domains support a phylogenetic affinity with A-domains from bacterial taxa and more rarely other fungi, a pattern most consistent with HGT. Our HGT survey comprised an extensive search of reference genomes across the tree of life available in NCBI RefSeq, as well as all of the gene models predicted for the Mucoromyocta and Zoopagomycota genomes. We find that 3% to 5% of all predicted gene models present in *Basidiobolous* genomes are consistent with signatures of HGT from bacteria. However, the percentages radically change to 41% to 66% of predicted SM core gene models with bacterial HGT evidence. These SM gene models with bacterial signatures are highly abundant, with NRPS and NRPS-like genes comprising the top 25 ontology categories of HGT genes in *B. meristosporus* (Supplementary Table 5). These results are consistent with the life history of *Basidiobolus*, where the fungus lives in close proximity with other microorganisms associated with the amphibian gut environment.

The additional analyses to discover distinct genetic features for HGT candidates showed that the largest differences found are in intron number and intron length, but not in nucleotide composition or by codon usage. A significantly smaller number of introns and smaller normalized intron length in HGT candidates provide more support to the HGT hypothesis, where we expected that bacterially transferred genes would maintain a smaller number and length of introns. Intronic expansion of transferred genes into fungal species after an HGT event appears to be rapid in order to reflect the genetic makeup across the genome (Da Lage et al. 2013) and can explain the introns in some of the HGT candidates. However, up to 60% of the HGT candidates still maintain absence of introns as expected for genes of bacterial origin. Finally, the nucleotide composition of HGT candidates and fungal genes were indistinguishable. Reports show that foreign genes with similar codon usage are more likely to become fixed on the receiving genome (Medrano-Soto et al. 2004, Amorós-Montoya et al. 2010, Tuller 2011). We interpet these results to indicate that horizontally transfered genes are evolving towards a similar nucleotide composition of the fungal genome based on the 5-mer/codon usage assay, but still maintain high protein similarity to and group with donor lineage copy in phylogenetic reconstructions.

The taxonomic survey of our HGT analysis shows that a diverse array of bacteria may have consistently contributed genetic information into the *Basidiobolus* genomes (Figure 6 and Supplementary Table 4). The most abundant bacterial taxonomic groups associated with HGT are the Proteobacteria, Firmicutes, Actinobacteria/high GC gram positive bacteria, and Bacteriodetes (Figure 6, Supplementary Table 4). The proportion of HGT for each taxonomic group appears to be consistent among the three *Basidiobolus* genomes, and there are consistencies between the most common taxonomic groups responsible for HGT in *Basidiobolus* and the reported composition of bacteria associated with the gut microbiome in amphibians (Bletz et al. 2016, Kohl et al. 2013) and reptiles (Colston and Jackson, 2016, Costello et al. 2010).

## Conclusions

Our results confirm that the majority of zygomycete fungi classified in Mucoromycota and Zoopagomycota do not possess a large genomic potential for secondary metabolism. Significant departures from this pattern exist, however, as exemplified by *Basidiobolus*, a genus with a complex genomic evolution and potential for considerable and diverse secondary metabolite production. First, it possesses larger than average genome with less than 8% content of repetitive regions, but a genetic plasticity to integrate and express extrinsic DNA. Second, the incorporation of extrinsic DNA is consistent with selection for increased SM production, especially gene models that are related to the capture of resources available in anaerobic conditions (iron chelation by siderophores) and metabolites that may play roles in antibiosis (surfactin-like genes) or host interaction. Third, the amphibian gut environment predisposes *Basidiobolus* to the acquisition of these SM core genes via HGT from co-inhabiting bacterial species. More information is needed to further test these hypotheses, including sequencing of additional *Basidiobolus* species with long read technologies; more accurate characterization of amphibian microbiomes that test positive for *Basidiobolus*; and LC-MSMS characterization of the *Basidiobolus* metabolome.

## Materials and Methods

### Data collection

Annotated genome and amino-acid translation of predicted gene model sequences for three isolates of two species within the genus *Basidiobolus* were used in this study: *Basidiobolus meristosporus* CBS 931.73 (Mondo et al. 2017), isolated from gecko dung in the locality of Lamco, Ivory Coast; *B. meristosporus* B9252 (Chibucos et al. 2016) isolated from human eye in Saudi Arabia; and *B. heterosporus* B8920 (Chibucos et al. 2016) isolated from plant debris in India. The genomic sequence of *B. meristosporus* CBS 931.73 was sequenced with PacBio and annotated by Mondo et al. (2017) and obtained from the US Department of Energy Joint Genome Institute *MycoCosm* genome portal (https://mycocosm.jgi.doe.gov; Grigoriev et al. 2014). Genomic sequences and annotation of *B. meristosporus* B9252 and *B. heterosporus* B8920 sequenced by Chibucos et al. (2016) were obtained directly from the authors. The raw reads for these two species are available in GenBank (Accession numbers GCA_000697375.1 and GCA_000697455.1). Data sources for the remaining genomes included in these analyses are available in Supplementary Table 1.

### Secondary metabolite gene cluster prediction

Secondary Metabolite gene clusters for *Basidiobolus* species were predicted with AntiSMASH v4.2.0 (Weber et al. 2017) and the Secondary Metabolite Unique Regions Finder (SMURF; Khaldi et al. 2010) from the annotated genomes of the three isolates used in this study. The AntiSMASH prediction was performed on local HPC, while SMURF predictions were obtained by submission of the genomes the SMURF web server (http://smurf.jcvi.org/). Predictions were contrasted manually to determine shared clusters of secondary metabolites. Predicted SM proteins were retrieved for the genomes of 66 additional Mucoromycota and Zoopagomycota species based on the SMURF predictions available at *MycoCosm* and by local prediction with AntiSMASH. Orthologous sets of core SM genes across *Basidiobolus* were identified using OrthoFinder (Emms et al. 2015).

### Secondary metabolite expression analysis

To assess the expression of predicted SM proteins in *B. meristosporus*, we calculated summarized counts of RNA transcript per million (TPM) for genes in *B. meristosporus* CBS 931.73 isolate by aligning RNA-Seq reads to the assembled *B. meristosporus* CBS 931.73 genome with HiSat v2.1.0 (Kim et al. 2019). The aligned sequence reads were processed with HTS-seq (Anders et al. 2014) to generate the counts of overlapping reads found for each gene and the normalized TPM for the genes was calculated using the cpm function in edgeR (Robinson et al. 2010). A distribution of RNA-Seq read counts per gene was plotted using the ggplot2 package in the R statistical framework (R Core Team 2018).

### Phylogenomic analyses

Phylogenetic analyses were used to assess the evolutionary relationships of the NRPS, PKS, and terpene cyclase/synthase predicted proteins. For the NRPS genes, the adenylation domains (A-domains) were identified by hmmsearch from HMMer 3.0 suite (Eddy 2004), using the A-domain profile reported by Bushley and Turgeon (2010) as a reference profile HMM. The predicted A domains were extracted from the resulting HMMER table into a FASTA file using the esl-reformat program included in the HMMER suite. The predicted A-domains for all *Basidiobolus*, Mucoromycota and Zoopagomycota species were added to an A-domain amino-acid alignment reported by Bushley and Turgeon (2010) with MAFFT v7 (Katoh et al. 2017) (Additional file 1). This reference alignment contains A-domains from NRPS proteins from nine major subfamilies of fungal and bacterial NRPS proteins. The phylogenetic domain tree was constructed using a maximum likelihood approach implemented in RAxML v. 8.2.11 (Stamatakis et al. 2014) with the JTT amino acid substitution matrix, after model selection using the PROTGAMMAAUTO option, and 1000 bootstrap replicates (raxmlHPC-PTHREADS -T 12 -n NRPS -s infile.fasta -f a -x 12345 -p 12345 -m PROTCATJTT –N 1000). A graphical representation of the A-domains from the NRPS/NRPS-like core gene models (Fig. *χ*) was constructed by coloring the A-domain position in the core gene according to its phylogenetic origin using the ggplot2 R package (Wickham, 2016).

For the PKS genes, the KS domains of all predicted PKS proteins from the *Basidiobolus*, Mucoromycota and Zoopagomycota species were identified using hmmsearch, using the KS domain profile (PF001009) available in PFAM v31 (Finn et al. 2009). The predicted KS domains for all *Basidiobolus*, Mucoromycota and Zoopagomycota species were added to an existing KS domain amino-acid alignment reported by Kroken et al. (2003) using MAFFT v7 (Katoh et al. 2017) (Additional file 2). This existing alignment contained KS domains from PKS proteins from reduced and unreduced PKS from bacterial and fungal species. The Kroken et al. (2003) database was expanded by adding predicted KS domains from PKS proteins of additional published fungal genomes in order to include more fungal diversity in the dataset: Eight Ascomycota isolates (*Aspergillus nidulans* FGSC A4, *Beauveria bassiana* ARSEF 2860, *Capronia coronata* CBS 617.96, *Capronia semiimmersa* CBS 27337, *Cladophialophora bantiana* CBS 173.52, *Cochliobolus victoriae* FI3 v1.0, *Microsporum canis* CBS 113480, *Trichoderma atroviride* v2.0), nine Basidiomycota isolates (*Acaromyces ingoldii* MCA 4198 v1.0, *Fibroporia radiculosa* TFFH 294, *Fomitiporia mediterranea* v1.0, *Gloeophyllum trabeum* v1.0, *Gymnopus luxurians* v1.0, *Laccaria bicolor* v2.0, *Microbotryum lychnidis−dioicae* p1A1 Lamole, *Piloderma croceum* F 1598 v1.0, *Pisolithus tinctorius* Marx 270 v1.0), four Neocallimastigomycota isolates (*Anaeromyces robustus* v1.0, *Orpinomyces sp*., *Piromyces finnis* v3.0, *Piromyces sp. E2* v1.0) and one Chytridiomycota species (*Spizellomyces punctatus* DAOM BR117). The predicted PKS proteins from additional species were all obtained from the DOE-JGI *MycoCosm* genome portal by searching for all genes with “PKS” on the SM annotation from *MycoCosm*. The KS domains were identified for the subset of PKS proteins using a HMMER KS profile as mentioned above. A phylogenetic tree was reconstructed using maximum likelihood using all KS domains using similar parameters described in the NRPS step.

The terpene cyclase (TC) proteins were predicted in AntiSMASH for all 69 assembled genome sequences from *Basidiobolus*, Mucoromycota and Zoopagomycota. To include additional fungal TC proteins in the phylogenetic reconstruction, we identified TC proteins from 58 published genomes of Dikarya isolates (21 Basidiomycetes and 37 Ascomycetes) available in DOE-JGI *MycoCosm*, each from a different family (Supplementary Table 3). One isolate per family was randomly selected for the analysis. To screen for TC proteins in Dikarya, OrthoFinder was used to build orthologous clusters of genes between *Basidiobolus*, Mucoromycota and Zoopagomycota TC and the Dikarya proteome dataset. Dikarya proteins clustered within orthologous groups that contain TC of *Basidiobolus*, Mucoromycota and Zoopagomycota were considered valid TC orthologs and used in downstream analyses. Finally, we identified bacterial TC to evaluate the potential for HGT into *Basidiobolus*, Mucoromycota or Zoopagomycota species. Bacterial TC’s were identified by screening each protein from the RefSeq, release 87 (May 2018, O’Leary et al. 2016) FASTA dataset against a BLAST database (Altschul et al. 1997), one from each of the zygomycete orthologous groups identified by OrthoFinder in the previous step, for a total of four protein sequences in the database. The BLASTP program was used to perform the searches, using an e-value of 1e^-10^ and the BLOSUM62 substitution matrix. Positive matches from the BLASTP assay were used as bacterial TC for subsequent analyses. A multi-sequence alignment containing all predicted TC from *Basidiobolus*, Mucoromycota and Zoopagomycota species, the orthologous TC from additional fungal species, and the TC proteins from reference bacterial predicted gene models result of the BLASTP search was performed in MAFFT v7 using the G-INS-1 algorithm for progressive global alignment (Katoh et al. 2017) (Additional file 3). A phylogenetic tree was reconstructed using maximum likelihood in RAxML using similar parameters as mentioned before.

### Horizontal gene transfer in Basidiobolus *species*

Identification of genic regions with evidence for horizontal gene transfer (HGT) from bacteria was performed by searching all translated proteins from the predicted gene models of the *Basidiobolus* isolates against a custom BLAST protein database. This database included all amino-acid sequences available in the NCBI RefSeq proteomics database and all available amino-acid sequences for Mucoromycota and Zoopagomycota species at the DOE-JGI *MycoCosm* genome portal. The proteins were searched with BLASTP against the combined RefSeq/Mucoromycota/Zoopagomycota custom database, using an e-value cutoff of 1e^-10^ and the BLOSUM62 substitution matrix. A summary of the taxonomy identifier for all best hits was obtained by identifying whether the best hit was a Mucoromycota/Zoopagomycota protein, or a RefSeq protein. We used the rentrez package (Winter, 2017) in the R statistical framework to extract the top-ten taxonomic identifiers from the NCBI database when hits did not correspond to Mucoromycota/Zoopagomycota. Proteins that had no hits to a fungal protein and only hits to a bacterial protein were considered candidate HGT genes.

To increase the accuracy of the prediction of genes product of HGT, we tested whether HGT candidate genes showed signs of errant assembly or in silica incorporation into the *Basidiobolus* genomes. The mean genomic read coverage of each candidate gene was calculated and compared to the mean coverage of the scaffold harboring the HGT candidate gene for all three *Basidiobolus* isolates. A z-score was calculated in R to determine the number of standard deviations of the HGT candidate from the mean coverage of the harboring scaffold. All HGT candidates with standard deviations greater or less than two were removed from the analysis. The genomic reads were mapped to each reference genome using BWA (Li and Durbin, 2009). Coverage per HGT and harboring scaffold was estimated using samtools (Li et al. 2009). A summary plot of the proportion of genes with bacterial best hits was constructed using the ggplot2 package in R.

To identify differences in the GC content between HGT candidates and genes of fungal origin, a paired t-test was performed between the mean GC content for HGT candidates and fungal genes for the predicted genes of *Basidiobolus meristosporus* CBS 931.73 as this species has the genome with longest contigs and best assemblies overall. In addition, to identify if the HGT candidate genes had a reduced number of introns than the fungal genes, a summary of the number of introns and normalized intron length (intron length divided by gene model length) was performed in the GenomicRanges R package (Lawrence et al. 2013). A Kruskal-Wallis test was performed in R to identify significant differences in the number of introns and the normalized intron length for the HGT candidates and the fungal genes. A nucleotide composition analysis based on 5-mers and codon usage was performed to observe differences between candidate HGT genes and fungal genes for *Basidiobolus meristosporus* CBS 931.73. The 5-mer analysis was conducted in all predicted coding sequences from the annotated gene models using the oligonucleotideFrequency function of the Biostring R package (Pagès et al. 2019). Codon usage was estimated using the uco function of the seqinr R package (Charif and Lobry 2007). Both these indices were divided by gene length to normalize the nucleotide composition by the effect of gene length. Principal component analyses were performed in R for both 5-mer and codon usage analysis to compare the HGT candidates to the fungal candidate genes. Lastly, the putative functions of the genes with evidence for HGT were summarized using the functional annotations available in the gene format file (GFF) of *B. meristosporus* CBS 931.73.

### Measuring siderophore activity in Basidiobolus

To measure the siderophore activity (chelation of ferric ions) of *Basidiobolus*, an assay of detection of siderophore activity based on the universal chrome azurol S (CAS) assay (Andrews et al. 2016) was performed for the strain *Basidiobolus meristosporus* CBS 931.73. This colorimetric assay uses a complex of Fe(III) – CAS – DDAPS (Surfactant). When this complex is combined with acetate yeast agar (AY agar), it results in a greenish-blue color, where the color changes to yellow upon the removal of the iron. A total of 450 ml of AY agar (Andrews et al. 2016) was mixed with 50 ml of autoclaved 10X CAS assay (20 ml of 10 mM Fe(NO_3_)_3_, 40 ml of 10 mM chrome azurol S, and 100 ml of 10 mM DDAPS). A 10 cm layer of the AY-CAS media was poured in small petri dishes and cooled. An upper layer of 10 cm of AY agar was poured after cooling, and the media was left to diffuse overnight and stored at 4°C for 24 hours. *B. meristosporus, Conidiobolus thromboides* FSU 785 (negative control), and *Cladosporium sp*. from the *herbarum* species complex PE-07 (positive control) were transferred into the AY-CAS plates by transferring a small amount of mycelium via a sterile toothpick and piercing the media in the center. The assay was performed in triplicate for each isolate used. Cultures were grown at room temperature for 12 days. Siderophore activity was measured as the area of the plate that has changed to yellow color, when compared to a negative control and an empty petri dish with AY-CAS agar. Pictures of the plates were taken. These images were imported into Adobe Photoshop CC 2019 and size-corrected to 5.5 cm (diameter of the small petri dish). All images were concatenated in the same file. The Color Range tool of Adobe Photoshop CC tool was used to measure the yellow area for all plates. The area measurements were exported into R, where an analysis of variance was performed to determine significant differences across the siderophore activity of the strains.

## Data availability

All additional files included as supplementary materials in this manuscript and custom scripts used for this analysis can be found in the ZyGoLife GitHub repository (https://github.com/zygolife/Basidiobolus_SM_repo).

## Acknowledgments

We would like to thank Dr. Vincent Bruno and Dr. Ashraf S. Ibrahim for kindly sharing the files of the genome annotation, assembly and gene predictions of *B. meristosporus* B9252 and *B. heterosporus* B8920; Dr. Marc Cubeta and Dr. Gregory Bonito for discussions and recommendations for additional experiments regarding siderophore assays and surfactin-like genes; Dr. Gregory Bonito for allowing us to use the unpublished genome sequences of *Mortierella humilis* PMI_1414, *Mortierella sp*. GBAus27b; Dr. Olafur S. Andrésson for allowing us to use the unpublished genome sequences of *Lobaria pulmonaria*; Dr. Paul Dryer for allowing us to use the unpublished genome sequence of *χanthoria parietina 46-1-SA22*; Dr. Francis Martin and Dr. Pierre Gladieux for allowing us to use the unpublished genome sequence of *Achaetomium strumarium CBS333*.*67*; Dr. Francis Martin for allowing us to use the unpublished genome sequences of *Boletus edulis V1* and *Amylostereum chailletii DWAch2*; Dr. Marie-Noëlle Rosso for allowing us to use the unpublished genome sequences of *Abortiporus biennis CIRM-BRFM1778;* Dr. Francis Martin and Dr. Gregory Bonito for allowing us to use the unpublished genome sequence of *Atractiellales rhizophila v2*.*0;* the 1000 Fungal Genomes Project for the use of the dikarya unpublished genomes; and Sabrina Heitmann, Dr. Ed Barge, Dr. Devin Leopold and Dr. Posy Busby for the *Cladosporium sp*. from the *herbarum* species complex PE-07 sample. This work was supported by the National Science Foundation (DEB-1441604 to JWS; DEB-1557110 and DEB-1441715 to JES). The work conducted by the U.S. Department of Energy Joint Genome Institute, a DOE Office of Science User Facility, is supported by the Office of Science of the U.S. Department of Energy under Contract No. DE-AC02-05CH11231. Any opinions, findings, and conclusions or recommendations expressed in this material are those of the author(s) and do not necessarily reflect the views of the National Science Foundation.

