## Supplementary Tables for "Phylogenomic analyses of non-Dikarya fungi supports horizontal gene transfer driving diversification of secondary metabolism in the amphibian gastrointestinal symbiont, *Basidiobolus*"

**Supplementary Table 1.** Isolates used in this study and respective secondary metabolite (SM) cluster prediction. All isolate genomes are available at the JGI MycoCosm data portal, except for *Basidiobolus meristosporus* B9252 and *Basidiobolus heterosporus* B8920 that were obtained from Chibucos et al. 2016. MycoCosm data for individual genomes can be accessed by appending Portal Id to ‘https://mycocosm.jgi.doe.gov/<Portal Id>”. NRPS: Non-ribosomal peptide synthetases. SM: Secondary metabolite core gene number PKS: Polyketide synthases, TC: Terpene Cyclases

| **Isolate/Species** | **Phylum** | **Subphylum** | **Sample identifier** | **Genome size (Mb)** | **Number of gene models** | **SM** | **HYBRID** | **NRPS** | **NRPS-Like** | **PKS** | **PKS-Like** | **TC** | **Mycocosm Portal ID /NCBI accesion** | **References** |
| --- | --- | --- | --- | --- | --- | --- | --- | --- | --- | --- | --- | --- | --- | --- |
| *Basidiobolus meristosporus B9252* | Zoopagomycota | Entomophthoromycotina | N161 | 103.9 | 13273 | 40 | 0 | 13 | 14 | 2 | 5 | 6 | GCA_000697375.1 | Chibucos et al. (2016) |
| *Basidiobolus heterosporus B8920* | Zoopagomycota | Entomophthoromycotina | N168 | 47.63 | 9331 | 23 | 0 | 10 | 8 | 1 | 0 | 4 | GCA_000697455.1 | Chibucos et al. (2016) |
| *Basidiobolus meristosporus CBS 931.73* | Zoopagomycota | Entomophthoromycotina | Basme2finSC | 89.49 | 16111 | 43 | 1 | 19 | 9 | 3 | 1 | 10 | Basme2finSC | Mondo et at. (2017) |
| *Conidiobolus coronatus NRRL28638 v1.0* | Zoopagomycota | Entomophthoromycotina | Conco1 | 39.9 | 4790 | 13 | 0 | 1 | 7 | 0 | 3 | 2 | Conco1 | Chang et al. (2015) |
| *Conidiobolus thromboides FSU 785 v1.0* | Zoopagomycota | Entomophthoromycotina | Conth1 | 24.64 | 4266 | 8 | 0 | 0 | 4 | 1 | 1 | 2 | Conth1 | Unpublished |
| *Coemansia mojavensis RSA 71 v1.0* | Zoopagomycota | Kickxellomycotina | Coemoj1 | 14.29 | 6859 | 11 | 0 | 1 | 0 | 6 | 1 | 3 | Coemoj1 | Unpublished |
| *Coemansia reversa NRRL 1564 v1.0* | Zoopagomycota | Kickxellomycotina | Coere1 | 21.84 | 4069 | 11 | 0 | 1 | 0 | 6 | 2 | 2 | Coere1 | Chang et al. (2015) |
| *Coemansia spiralis RSA 1278 v1.0* | Zoopagomycota | Kickxellomycotina | Coespi1 | 18.93 | 4298 | 15 | 0 | 1 | 0 | 11 | 1 | 2 | Coespi1 | Unpublished |
| *Dimargaris cristalligena RSA 468 single-cell v1.0* | Zoopagomycota | Kickxellomycotina | DimcrSC1 | 30.78 | 3721 | 33 | 0 | 21 | 7 | 0 | 2 | 3 | DimcrSC1 | Ahrendt et al. (2018) |
| *Kickxella alabastrina RSA 675 v1.0* | Zoopagomycota | Kickxellomycotina | Kicala1 | 22.26 | 4190 | 7 | 0 | 0 | 1 | 2 | 2 | 2 | Kicala1 | Unpublished |
| *Linderina pennispora ATCC 12442 v1.0* | Zoopagomycota | Kickxellomycotina | Linpe1 | 26.2 | 4785 | 23 | 0 | 1 | 0 | 15 | 3 | 4 | Linpe1 | Mondo et at. (2017) |
| *Martensiomyces pterosporus CBS 209.56 v1.0* | Zoopagomycota | Kickxellomycotina | Marpt1 | 19.82 | 4577 | 23 | 0 | 1 | 0 | 5 | 14 | 3 | Marpt1 | Unpublished |
| *Ramicandelaber brevisporus CBS 109374 v1.0* | Zoopagomycota | Kickxellomycotina | Rambr1 | 25.53 | 5177 | 5 | 0 | 0 | 1 | 1 | 1 | 2 | Rambr1 | Unpublished |
| *Smittium culicis ID-206-W2* | Zoopagomycota | Kickxellomycotina | SmiculW2_1 | 71.05 | 5686 | 4 | 0 | 0 | 1 | 2 | 1 | NA | SmiculW2_1 | Wang et al. (2017) |
| *Smittium culicis GSMNP* | Zoopagomycota | Kickxellomycotina | SmicuMNP_2 | 77.12 | 6440 | 4 | 0 | 0 | 1 | 2 | 1 | NA | SmicuMNP_2 | Wang et al. (2017) |
| *Smittium mucronatum ALG-7-W6* | Zoopagomycota | Kickxellomycotina | Smimuc2 | 102.35 | 4347 | 5 | 0 | 0 | 1 | 3 | 1 | NA | Smimuc2 | Wang et al. (2017) |
| *Zancudomyces culisetae COL-18-3* | Zoopagomycota | Kickxellomycotina | Zancul2 | 28.64 | 4222 | 5 | 0 | 0 | 1 | 0 | 3 | 1 | Zancul2 | Wang et al. (2017) |
| *Piptocephalis cylindrospora RSA 2659 single-cell v3.0* | Zoopagomycota | Zoopagomycotina | Pipcy3_1 | 10.75 | 2143 | 3 | 0 | 0 | 0 | 0 | 1 | 2 | Pipcy3_1 | Ahrendt et al. (2018) |
| *Syncephalis fuscata S228 v1.0* | Zoopagomycota | Zoopagomycotina | Synfus1 | 29.36 | 4726 | 5 | 0 | 1 | 0 | 0 | 1 | 3 | Synfus1 | Unpublished |
| *Syncephalis plumigaleata NRRL S24 v1.0* | Zoopagomycota | Zoopagomycotina | Synplu1 | 32.91 | 8130 | 4 | 0 | 1 | 0 | 0 | 1 | 2 | Synplu1 | Unpublished |
| *Syncephalis pseudoplumigaleata Benny S71-1 single-cell v1.0* | Zoopagomycota | Zoopagomycotina | Synps1 | 16.27 | 2941 | 6 | 0 | 1 | 0 | 0 | 1 | 4 | Synps1 | Ahrendt et al. (2018) |
| *Thamnocephalis sphaerospora RSA 1356 single-cell v1.0* | Zoopagomycota | Zoopagomycotina | Thasp1 | 18.2 | 3697 | 4 | 0 | 0 | 1 | 0 | 1 | 2 | Thasp1 | Ahrendt et al. (2018) |
| *Rhizophagus irregularis DAOM 181602 v1.0* | Mucoromycota | Glomeromycotina | Gloin1 | 91.08 | 10868 | 7 | 0 | 1 | 0 | 0 | 1 | 5 | Gloin1 | Tisserant et al. (2013) |
| *Rhizophagus irregularis DAOM 197198 v2.0* | Mucoromycota | Glomeromycotina | Rhiir2_1 | 136.81 | 11110 | 7 | 0 | 1 | 1 | 0 | 1 | 4 | Rhiir2_1 | Chen et al. (2018) |
| *Rhizophagus irregularis A1 v1.0* | Mucoromycota | Glomeromycotina | RhiirA1_1 | 125.87 | 11046 | 4 | 0 | 1 | 1 | 0 | 2 | NA | RhiirA1_1 | Chen et al. (2018) |
| *Rhizophagus irregularis A4 v1.0* | Mucoromycota | Glomeromycotina | RhiirA4 | 138.3 | 10848 | 3 | 0 | 1 | 0 | 0 | 2 | NA | RhiirA4 | Chen et al. (2018) |
| *Rhizophagus irregularis A5 v1.0* | Mucoromycota | Glomeromycotina | RhiirA5 | 131.46 | 10893 | 2 | 0 | 1 | 0 | 0 | 1 | NA | RhiirA5 | Chen et al. (2018) |
| *Rhizophagus irregularis B3 v1.0* | Mucoromycota | Glomeromycotina | RhiirB3 | 124.89 | 10707 | 3 | 0 | 1 | 0 | 0 | 2 | NA | RhiirB3 | Chen et al. (2018) |
| *Rhizophagus irregularis C2 v1.0* | Mucoromycota | Glomeromycotina | RhiirC2 | 122.97 | 10677 | 4 | 0 | 1 | 1 | 0 | 2 | NA | RhiirC2 | Chen et al. (2018) |
| *Lobosporangium transversale NRRL 3116 v1.0* | Mucoromycota | Morteriellomycotina | Lobtra1 | 42.77 | 7348 | 6 | 0 | 1 | 0 | 1 | 1 | 3 | Lobtra1 | Mondo et at. (2017) |
| *Mortierella elongata AG-77 v2.0* | Mucoromycota | Morteriellomycotina | Morel2 | 49.86 | 9032 | 6 | 0 | 0 | 3 | 0 | 1 | 2 | Morel2 | Uehling et al. (2017) |
| *Mortierella multidivaricata RSA 2152 T v1.0* | Mucoromycota | Morteriellomycotina | Mormul1 | 37.67 | 7462 | 7 | 0 | 0 | 3 | 1 | 1 | 2 | Mormul1 | Unpublished |
| *Mortierella verticillata NRRL 6337* | Mucoromycota | Morteriellomycotina | Morve1 | 41.85 | 8493 | 8 | 0 | 4 | 2 | 1 | 1 | NA | Morve1 | Broad Institute |
| *Mortierella sp. GBAus27b* | Mucoromycota | Morteriellomycotina | MorGBAus27b_1 | 44.97 | 13953 | 21 | 0 | 3 | 13 | 1 | 1 | 3 | MorGBAus27b_1 | Unpublished (Gregory Bonito) |
| *Absidia repens NRRL 1336 v1.0* | Mucoromycota | Mucoromycotina | Absrep1 | 47.42 | 8575 | 14 | 0 | 1 | 1 | 1 | 1 | 10 | Absrep1 | Mondo et at. (2017) |
| *Backusella circina FSU 941 v1.0* | Mucoromycota | Mucoromycotina | Bacci1 | 48.65 | 9250 | 16 | 0 | 1 | 5 | 2 | 1 | 7 | Bacci1 | Unpublished |
| *Blakeslea trispora NRRL 2456 v1.0* | Mucoromycota | Mucoromycotina | Blatri1 | 37.51 | 6062 | 12 | 0 | 1 | 3 | 1 | 1 | 6 | Blatri1 | Unpublished |
| *Absidia padenii NRRL 2977 v1.0* | Mucoromycota | Mucoromycotina | Chlpad1 | 34.33 | 7616 | 9 | 0 | 0 | 0 | 0 | 0 | 9 | Chlpad1 | Unpublished |
| *Circinella umbellata NRRL1351 v1.0* | Mucoromycota | Mucoromycotina | Cirumb1 | 51.27 | 7773 | 12 | 0 | 0 | 4 | 1 | 1 | 6 | Cirumb1 | Unpublished |
| *Cokeromyces recurvatus NRRL 2243 v1.0* | Mucoromycota | Mucoromycotina | Cokrec1 | 28.12 | 5928 | 13 | 0 | 1 | 2 | 2 | 1 | 7 | Cokrec1 | Unpublished |
| *Cunninghamella echinulata NRRL 1382 v1.0* | Mucoromycota | Mucoromycotina | Cunech1 | 29 | 5915 | 10 | 0 | 1 | 0 | 1 | 1 | 7 | Cunech1 | Unpublished |
| *Dichotomocladium elegans RSA 919- v1.0* | Mucoromycota | Mucoromycotina | Dicele1 | 39.78 | 6567 | 8 | 0 | 0 | 1 | 1 | 1 | 5 | Dicele1 | Unpublished |
| *Fennellomyces sp. T-0311 v1.0* | Mucoromycota | Mucoromycotina | Fenlin1 | 45.9 | 7672 | 14 | 0 | 0 | 5 | 1 | 1 | 7 | Fenlin1 | Unpublished |
| *Gilbertella persicaria var. persicaria CBS 190.32-T v1.0* | Mucoromycota | Mucoromycotina | Gilper1 | 25.73 | 6150 | 12 | 0 | 1 | 3 | 1 | 1 | 6 | Gilper1 | Unpublished |
| *Gongronella butleri v1.0* | Mucoromycota | Mucoromycotina | Gonbut1 | 33.01 | 6298 | 12 | 0 | 1 | 2 | 1 | 1 | 7 | Gonbut1 | Unpublished |
| *Hesseltinella vesiculosa NRRL3301 v2.0* | Mucoromycota | Mucoromycotina | Hesve2finisherSC | 27.22 | 6117 | 11 | 0 | 1 | 2 | 1 | 1 | 6 | Hesve2finisherSC | Mondo et at. (2017) |
| *Mucor cordense RSA 1222 v1.0* | Mucoromycota | Mucoromycotina | Kircor1 | 40.95 | 9104 | 22 | 0 | 1 | 7 | 2 | 1 | 11 | Kircor1 | Unpublished |
| *Lichtheimia corymbifera JMRC:FSU:9682* | Mucoromycota | Mucoromycotina | Liccor1 | 33.53 | 7358 | 13 | 0 | 0 | 5 | 1 | 1 | 6 | Liccor1 | Schwartze et al. (2014) |
| *Lichtheimia hyalospora v1.0* | Mucoromycota | Mucoromycotina | Lichy1 | 33.28 | 6757 | 15 | 0 | 0 | 6 | 1 | 1 | 7 | Lichy1 | Unpublished |
| *Mucor circinelloides CBS277.49 v2.0* | Mucoromycota | Mucoromycotina | Mucci2 | 36.6 | 11719 | 15 | 0 | 1 | 4 | 2 | 1 | 7 | Mucci2 | Corrochano et al. (2016) |
| *Mycotypha africana NRRL 2978 v1.0* | Mucoromycota | Mucoromycotina | Mycafr1 | 29.2 | 5937 | 11 | 0 | 1 | 1 | 2 | 1 | 6 | Mycafr1 | Unpublished |
| *Parasitella parasitica v1.0* | Mucoromycota | Mucoromycotina | Parpar1 | 32.77 | 6765 | 13 | 0 | 1 | 2 | 2 | 1 | 7 | Parpar1 | Unpublished |
| *Phascolomyces articulosus v1.0* | Mucoromycota | Mucoromycotina | Phaart1 | 47.61 | 7974 | 14 | 0 | 0 | 7 | 1 | 1 | 5 | Phaart1 | Unpublished |
| *Pilaira anomala RSA1997 v1.0* | Mucoromycota | Mucoromycotina | Pilano1 | 34.9 | 6925 | 13 | 0 | 1 | 3 | 1 | 1 | 7 | Pilano1 | Unpublished |
| *Pilobolus umbonatus NRRL 6349 v1.0* | Mucoromycota | Mucoromycotina | Pilumb1 | 34.77 | 5872 | 8 | 0 | 1 | 0 | 1 | 1 | 5 | Pilumb1 | Unpublished |
| *Radiomyces spectabilis NRRL 2753 v1.0* | Mucoromycota | Mucoromycotina | Radspe1 | 30.41 | 6228 | 13 | 0 | 0 | 3 | 1 | 1 | 8 | Radspe1 | Unpublished |
| *Rhizopus microsporus var. chinensis CCTCC M201021* | Mucoromycota | Mucoromycotina | Rhich1 | 45.74 | 10264 | 22 | 0 | 2 | 6 | 1 | 2 | 11 | Rhich1 | Wang et al. (2013) |
| *Rhizopus microsporus ATCC11559 v1.0* | Mucoromycota | Mucoromycotina | Rhimi_ATCC11559_1 | 24.08 | 5856 | 12 | 0 | 1 | 3 | 1 | 1 | 6 | Rhimi_ATCC11559_1 | Lastovetsky et al. (2016) |
| *Rhizopus microsporus var. microsporus ATCC52814 v1.0* | Mucoromycota | Mucoromycotina | Rhimi_ATCC52814_1 | 24.95 | 5849 | 13 | 0 | 1 | 3 | 1 | 1 | 7 | Rhimi_ATCC52814_1 | Lastovetsky et al. (2016) |
| *Rhizopus microsporus var. microsporus ATCC52813 v1.0* | Mucoromycota | Mucoromycotina | Rhimi1_1 | 25.97 | 5811 | 12 | 0 | 1 | 3 | 1 | 1 | 6 | Rhimi1_1 | Mondo et at. (2017) |
| *Saksenaea vasiformis B4078* | Mucoromycota | Mucoromycotina | Sakvas1 | 42.5 | 5877 | 9 | 0 | 1 | 1 | 1 | 0 | 6 | Sakvas1 | Chibucos et al. (2016) |
| *Spinellus fusiger NRRL 22323 v1.0* | Mucoromycota | Mucoromycotina | Spifus1 | 38.2 | 5394 | 14 | 0 | 0 | 6 | 1 | 1 | 6 | Spifus1 | Unpublished |
| *Sporodiniella umbellata MES 1446 v1.0* | Mucoromycota | Mucoromycotina | Spoumb1 | 26.24 | 5768 | 15 | 0 | 1 | 6 | 1 | 1 | 6 | Spoumb1 | Unpublished |
| *Syncephalastrum racemosum:13706* | Mucoromycota | Mucoromycotina | Synrac1 | 30.75 | 11124 | 15 | 0 | 0 | 7 | 1 | 1 | 6 | Synrac1 | Mondo et at. (2017) |
| *Mucor heterogamus NRRL 1489 v1.0* | Mucoromycota | Mucoromycotina | Zyghet1 | 54.49 | 14998 | 18 | 0 | 2 | 3 | 1 | 1 | 11 | Zyghet1 | Unpublished |
| *Mortierella humilis PMI_1414* | Mucoromycota | Mucoromycotina | Morhum1 | 36.2 | 12012 | 10 | 0 | 3 | 2 | 1 | 1 | 3 | Morhum1 | Unpublished (Gregory Bonito) |
| *Rhizopus delemar 99-880 from Broad* | Mucoromycota | Mucoromycotina | Rhior3 | 46.1 | 17467 | 10 | 0 | 1 | 6 | 1 | 1 | 1 | Rhior3 | Ma et al. (2009) |
| *Phycomyces blakesleeanus NRRL1555 v2.0* | Mucoromycota | Mucoromycotina | Phybl2 | 53.9 | 16528 | 12 | 0 | 1 | 2 | 1 | 1 | 7 | Phybl2 | Corrochano et al. (2016) |
| *Umbelopsis ramanniana AG # v1.0* | Mucoromycota | Mucoromycotina | Umbra1 | 23.08 | 9931 | 12 | 0 | 1 | 4 | 0 | 1 | 6 | Umbra1 | Unpublished |

**Supplementary Table 2.** Number of SM gene clusters identified for the three isolates of *Basidiobolus* analyzed in this report

| **Genome sequence** | **Gene model** | **SM class** | **Cluster number** | **Additional genes predicted in cluster (Protein ID)** |
| --- | --- | --- | --- | --- |
| *B .meristosporus* B9252 | N161.mRNA.1413.1 | Terpene | 1 | N161_02321;N161_02322;N161_02323;N161_02324;N161_02325;N161_02326 |
| *B .meristosporus* B9252 | N161.mRNA.1431.1 | NRPS | 2 | N161_00522;N161_00523;N161_00524 |
| *B .meristosporus* B9252 | N161.mRNA.1433.1 | NRPS-Like | 2 | N161_00522;N161_00523;N161_00524 |
| *B .meristosporus* B9252 | N161.mRNA.1485.1 | NRPS | 3 | N161_03744;N161_03745 |
| *B .meristosporus* B9252 | N161.mRNA.1486.1 | NRPS-Like | 3 | N161_03744;N161_03745 |
| *B .meristosporus* B9252 | N161.mRNA.2152.1 | NRPS | 4 | N161_10119 |
| *B .meristosporus* B9252 | N161.mRNA.3482.1 | NRPS-Like | 5 | N161_01570;N161_01571;N161_01572;N161_01573;N161_01574;N161_01575;N161_01576;N161_01577;N161_01578;N161_01579;N161_01580;N161_01581 |
| *B .meristosporus* B9252 | N161.mRNA.4115.1 | NRPS | 6 | N161_10439;N161_10440;N161_10441 |
| *B .meristosporus* B9252 | N161.mRNA.4211.1 | NRPS | 7 | N161_02960 |
| *B .meristosporus* B9252 | N161.mRNA.4324.1 | NRPS | 8 | N161_04345;N161_04346;N161_04347;N161_04348;N161_04349;N161_04350 |
| *B .meristosporus* B9252 | N161.mRNA.4477.1 | NRPS | 9 | N161_03649 |
| *B .meristosporus* B9252 | N161.mRNA.5264.1 | Terpene | 10 | N161_03756;N161_03757;N161_03758;N161_03759;N161_03760;N161_03761;N161_03762 |
| *B .meristosporus* B9252 | N161.mRNA.5262.1 | Terpene | 10 | N161_03756;N161_03757;N161_03758;N161_03759;N161_03760;N161_03761;N161_03762 |
| *B .meristosporus* B9252 | N161.mRNA.5494.1 | NRPS-Like | 11 | N161_11865;N161_11866;N161_11867 |
| *B .meristosporus* B9252 | N161.mRNA.5537.1 | NRPS-Like | 12 | N161_01252;N161_01253 |
| *B .meristosporus* B9252 | N161.mRNA.5981.1 | NRPS-Like | 13 | N161_06505;N161_06506;N161_06507;N161_06508 |
| *B .meristosporus* B9252 | N161.mRNA.6146.1 | PKS | 14 | N161_03348;N161_03349;N161_03350;N161_03351;N161_03352;N161_03353 |
| *B .meristosporus* B9252 | N161.mRNA.6846.1 | NRPS-Like | 15 | N161_00937;N161_00938;N161_00939;N161_00940;N161_00941;N161_00942;N161_00943;N161_00944;N161_00945 |
| *B .meristosporus* B9252 | N161.mRNA.6935.1 | NRPS-Like | 16 | N161_02910 |
| *B .meristosporus* B9252 | N161.mRNA.7156.1 | NRPS-Like | 17 | N161_07755 |
| *B .meristosporus* B9252 | N161.mRNA.8304.1 | NRPS | 18 | N161_00029;N161_00030;N161_00031;N161_00032;N161_00033;N161_00034 |
| *B .meristosporus* B9252 | N161.mRNA.8432.1 | NRPS | 19 | N161_08819;N161_08820;N161_08821;N161_08822;N161_08823;N161_08824;N161_08825;N161_08826;N161_08827 |
| *B .meristosporus* B9252 | N161.mRNA.8434.1 | PKS-Like | 19 | N161_08819;N161_08820;N161_08821;N161_08822;N161_08823;N161_08824;N161_08825;N161_08826;N161_08827 |
| *B .meristosporus* B9252 | N161.mRNA.9145.1 | NRPS | 20 | N161_05517 |
| *B .meristosporus* B9252 | N161.mRNA.9317.1 | NRPS | 21 | N161_10084 |
| *B .meristosporus* B9252 | N161.mRNA.9497.1 | NRPS-Like | 22 | N161_07963;N161_07964;N161_07965;N161_07966;N161_07967;N161_07968 |
| *B .meristosporus* B9252 | N161.mRNA.9639.1 | NRPS | 23 | N161_02356 |
| *B .meristosporus* B9252 | N161.mRNA.10391.1 | PKS | 24 | N161_01106;N161_01107 |
| *B .meristosporus* B9252 | N161.mRNA.11155.1 | NRPS-Like | 25 | N161_04188;N161_04189;N161_04190;N161_04191;N161_04192;N161_04193;N161_04194;N161_04195;N161_04196;N161_04197;N161_04198;N161_04199;N161_04200;N161_04201 |
| *B .meristosporus* B9252 | N161.mRNA.11155.1 | NRPS-Like | 26 | N161_04197 |
| *B .meristosporus* B9252 | N161.mRNA.11289.1 | NRPS | 27 | N161_01308 |
| *B .meristosporus* B9252 | N161.mRNA.12214.1 | Terpene | 28 | N161_10522;N161_10523;N161_10524;N161_10525;N161_10526;N161_10527 |
| *B .meristosporus* B9252 | N161.mRNA.13295.1 | NRPS-Like | 29 | N161_06649;N161_06650;N161_06651;N161_06652 |
| *B .meristosporus* B9252 | N161.mRNA.13969.1 | Terpene | 30 | N161_00991;N161_00992 |
| *B .meristosporus* B9252 | N161.mRNA.14278.1 | HYBRID | 31 | N161_09308;N161_09309;N161_09310;N161_09311;N161_09312;N161_09313;N161_09314;N161_09315 |
| *B .meristosporus* B9252 | N161.mRNA.14365.1 | Terpene | 32 | N161_10409;N161_10410;N161_10411 |
| *B .meristosporus* B9252 | N161.mRNA.12007.1 | NRPS-Like | 33 | N161_04229;N161_04230;N161_04231;N161_04232;N161_04233;N161_04234;N161_04235;N161_04236 |
| *B .meristosporus* B9252 | N161.mRNA.5862.1 | NRPS-Like | 34 | N161_03051 |
| *B .meristosporus* B9252 | N161.mRNA.8699.1 | NRPS-Like | 35 | N161_10399;N161_07977;N161_11287;N161_04081;N161_07890;N161_00310;N161_02875;N161_03816;N161_03190;N161_12282;N161_05360;N161_02320;N161_06895;N161_09821 |
| *B .meristosporus* B9252 | N161.mRNA.8700.1 | NRPS-Like | 36 | N161_08819;N161_01810;N161_05857;N161_07278;N161_08079;N161_07843;N161_05690;N161_07209;N161_01891;N161_06243;N161_01644;N161_05769;N161_11037;N161_06895;N161_09821 |
| *B .meristosporus* B9252 | N161.mRNA.204.1 | PKS-Like | 37 | N161_12043 |
| *B .meristosporus* B9252 | N161.mRNA.4846.1 | PKS-Like | 38 | N161_09808 |
| *B .meristosporus* B9252 | N161.mRNA.9939.1 | PKS-Like | 39 | N161_09808 |
| *B .meristosporus* B9252 | N161.mRNA.9973.1 | PKS-Like | 40 | N161_02595 |
| *B .meristosporus* CBS 931.73 | 366903 | NRPS | 1 | 295637;366903;366910;295640;310315;5321 |
| *B .meristosporus* CBS 931.73 | 322666 | NRPS-Like | 2 | 322664;344892;404256;310870;344895;252485;322666;296369;322667;374951 |
| *B .meristosporus* CBS 931.73 | 328337 | Terpene | 3 | 334011;311465;328337;311467;276042 |
| *B .meristosporus* CBS 931.73 | 146993 | NRPS-Like | 4 | 382072;323047;323048;311608;311609;334162;404891;146993;345979;297308;276322;297310;297311 |
| *B .meristosporus* CBS 931.73 | 387529 | NRPS | 5 | 387529;312194;277549;277552;298096;277554;334759;298099 |
| *B .meristosporus* CBS 931.73 | 278430 | Terpene | 6 | 99487;278430;99499;347625;312651;323585;323586;323587;210480;256087;312655 |
| *B .meristosporus* CBS 931.73 | 323585 | Terpene | 6 | 99487;278430;99499;347625;312651;323585;323586;323587;210480;256087;312655 |
| *B .meristosporus* CBS 931.73 | 298977 | NRPS | 7 | 298977;312902;312903;328970;348020;348021 |
| *B .meristosporus* CBS 931.73 | 368581 | NRPS | 8 | 349684;300169;368581;215133;258492;313863;280838;406759 |
| *B .meristosporus* CBS 931.73 | 349800 | NRPS | 9 | 11775;349800;349801;349802;215367;313931;313932;313933;406809;368907;313935 |
| *B .meristosporus* CBS 931.73 | 300898 | NRPS-Like | 10 | 314493;217438;314495;19557;281917;259598;281923;300898;300899;300900;300901 |
| *B .meristosporus* CBS 931.73 | 300898 | NRPS-Like | 10 | 314493;217438;314495;19557;281917;259598;281923;300898;300899;300900;300901 |
| *B .meristosporus* CBS 931.73 | 282246 | Terpene | 11 | 301086;21234;21255;282246;314665;407354;218038;301092 |
| *B .meristosporus* CBS 931.73 | 301092 | Terpene | 11 | 301086;21234;21255;282246;314665;407354;218038;301092 |
| *B .meristosporus* CBS 931.73 | 260222 | Terpene | 12 | 22761;218744;218712;22802;260217;314835;351312;260222;337205;351315 |
| *B .meristosporus* CBS 931.73 | 301341 | Terpene | 13 | 314880;314882;301338;24046;337244;301341;282611;301343 |
| *B .meristosporus* CBS 931.73 | 372159 | NPRS | 14 | 301473;24895;314994;372159;219244;372167;314995;407608;314997;182058;219234;315001;351566;315003 |
| *B .meristosporus* CBS 931.73 | 351909 | NRPS | 15 | 372749;351909 |
| *B .meristosporus* CBS 931.73 | 372749 | NRPS | 15 | 372749;351909 |
| *B .meristosporus* CBS 931.73 | 337511 | NRPS-Like | 16 | 337511;301681;329810;301683;220047 |
| *B .meristosporus* CBS 931.73 | 372991 | NRPS | 17 | 372991;154765;139122;220322;301747;301748;407790;301750;337567;315257 |
| *B .meristosporus* CBS 931.73 | 373247 | NRPS | 18 | 373247;220753;28376;220738;329867;315346;315348;324755;315350;315351;315353 |
| *B .meristosporus* CBS 931.73 | 373940 | NRPS | 19 | 373940;302148;352631;31440;283912;352634 |
| *B .meristosporus* CBS 931.73 | 221915 | NRPS-Like | 20 | 221900;221945;352795;32535;337961;221950;32530;337963;32572;315660;315661;315662;221915;261780;337969;352803;374274;352805 |
| *B .meristosporus* CBS 931.73 | 375475 | NRPS | 21 | 375475 |
| *B .meristosporus* CBS 931.73 | 338397 | NRPS | 22 | 330193;302810;223759;316123;375580;338397;353622;302814;353623;353624 |
| *B .meristosporus* CBS 931.73 | 375580 | NRPS | 22 | 330193;302810;223759;316123;375580;338397;353622;302814;353623;353624 |
| *B .meristosporus* CBS 931.73 | 375613 | NRPS | 23 | 37789;316128;37852;223771;338402;37874;284951;375613;223806;353650;302826;353651;353652 |
| *B .meristosporus* CBS 931.73 | 377413 | NRPS | 24 | 303544;377413;408976;316765;45707;303549;316769;316770;263779 |
| *B .meristosporus* CBS 931.73 | 409012 | Terpene | 25 | 316823;286132;409012;316830 |
| *B .meristosporus* CBS 931.73 | 304520 | Terpene | 26 | 330193;302810;223759;316123;375580;338397;353622;302814;353623;353624 |
| *B .meristosporus* CBS 931.73 | 229390 | Terpene | 26 | 330193;302810;223759;316123;375580;338397;353622;302814;353623;353624 |
| *B .meristosporus* CBS 931.73 | 382467 | NRPS-Like | 27 | 318559;318560;318561;340521;318563;318564;305608;357897;318565;232920;382467;357901;66822 |
| *B .meristosporus* CBS 931.73 | 340613 | NRPS-Like | 28 | 340608;305718;156797;305719;326180;340610;318645;318646;233298;289283;340613;233269 |
| *B .meristosporus* CBS 931.73 | 237744 | PKS | 29 | 306991;319740;341570;306994;237745;237740;306997;237744;237746 |
| *B .meristosporus* CBS 931.73 | 291346 | NRPS-Like | 30 | 319840;291346;238215 |
| *B .meristosporus* CBS 931.73 | 320108 | NRPS-Like | 31 | 411566;291781;320108;269502 |
| *B .meristosporus* CBS 931.73 | 387714 | NRPS-Like | 32 | 152918;239552;291892;341975;320182;291896;341977;331710;387714;360885;320185;88353 |
| *B .meristosporus* CBS 931.73 | 307892 | NRPS | 33 | 307892;307893;241197;307895;320516;241198;307898 |
| *B .meristosporus* CBS 931.73 | 292783 | PKS | 34 | 292783;320693;331891;241971;308095;361867;411992;241981 |
| *B .meristosporus* CBS 931.73 | 343011 | NRPS | 35 | 309006;321539;343011 |
| *B .meristosporus* CBS 931.73 | 363930 | NRPS | 36 | 363925;363926;309098;309099;309100;309101;363930;363931 |
| *B .meristosporus* CBS 931.73 | 290138 | PKS | 37 | 290138 |
| *B .meristosporus* CBS 931.73 | 207695 | PKS-Like | 38 | 207695 |
| *B. heterosporus* B8920 | N168_02885 | NRPS | 1 | N168_02882;N168_02883;N168_02884;N168_02885;N168_02886;N168_02887;N168_02888;N168_02889;N168_02890 |
| *B. heterosporus* B8920 | N168_04958 | PKS | 2 | N168_04958;N168_04959;N168_04960;N168_04961;N168_04962;N168_04963;N168_04964;N168_04965 |
| *B. heterosporus* B8920 | N168_00992 | NRPS | 3 | N168_00992 |
| *B. heterosporus* B8920 | N168_00027 | Terpene | 4 | N168_05324 |
| *B. heterosporus* B8920 | N168_05324 | NRPS-Like | 5 | N168_00025;N168_00026;N168_00027;N168_00028;N168_00029 |
| *B. heterosporus* B8920 | N168_07733 | NRPS | 6 | N168_07733 |
| *B. heterosporus* B8920 | N168_08644 | Terpene | 7 | N168_08640;N168_08641;N168_08642;N168_08643;N168_08644;N168_08645;N168_08646;N168_08647 |
| *B. heterosporus* B8920 | N168_05934 | NRPS | 8 | N168_05932;N168_05933;N168_05934 |
| *B. heterosporus* B8920 | N168_07140 | NRPS | 9 | N168_07138;N168_07139;N168_07140 |
| *B. heterosporus* B8920 | N168_02949 | NRPS-Like | 10 | N168_02949;N168_02950;N168_02951 |
| *B. heterosporus* B8920 | N168_03036 | NRPS-Like | 11 | N168_03036 |
| *B. heterosporus* B8920 | N168_04239 | NRPS-Like | 12 | N168_04234;N168_04235;N168_04236;N168_04237;N168_04238;N168_04239 |
| *B. heterosporus* B8920 | N168_06479 | NRPS | 13 | N168_06474;N168_06475;N168_06476;N168_06477;N168_06478;N168_06479 |
| *B. heterosporus* B8920 | N168_00034 | NRPS | 14 | N168_00034;N168_00035 |
| *B. heterosporus* B8920 | N168_00176 | NRPS | 15 | N168_00168;N168_00169;N168_00170;N168_00171;N168_00172;N168_00173;N168_00174;N168_00175;N168_00176 |
| *B. heterosporus* B8920 | N168_04712 | NRPS-Like | 16 | N168_04712 |
| *B. heterosporus* B8920 | N168_00255 | NRPS-Like | 17 | N168_00250;N168_00251;N168_00252;N168_00253;N168_00254;N168_00255;N168_00256 |
| *B. heterosporus* B8920 | N168_08721 | NRPS | 18 | N168_08721 |
| *B. heterosporus* B8920 | N168_05966 | NRPS | 19 | N168_05966;N168_05967 |
| *B. heterosporus* B8920 | N168_04564 | Terpene | 20 | N168_04560;N168_04561;N168_04562;N168_04563;N168_04564;N168_04565;N168_04566 |
| *B. heterosporus* B8920 | N168_00768 | NRPS-Like | 21 | N168_00759;N168_00760;N168_00761;N168_00762;N168_00763;N168_00764;N168_00765;N168_00766;N168_00767;N168_00768;N168_00769;N168_00770 |
| *B. heterosporus* B8920 | N168_03279 | Terpene | 22 | N168_03276;N168_03277;N168_03278;N168_03279;N168_03280;N168_03281;N168_03282;N168_03283 |
| *B. heterosporus* B8920 | N168_08580 | NRPS-Like | 23 | N168_08580 |

**Supplementary Table 3.** Isolates of Basidiomycete and Ascomycete species used for terpene cyclase (TC) prediction in Dykaria. All isolate genomes are available at the JGI MycoCosm data portal. 1KFG: 1000 Fungal Genomes Program.

| **JGI Identifier** | **Phylum** | **Subphylum** | **Class** | **Order** | **Family** | **Species** | **Reference** |
| --- | --- | --- | --- | --- | --- | --- | --- |
| Aplpr1 | Ascomycota | Pezizomycotina | Dothideomycetes | Botryosphaeriales | Botryosphaeriaceae | *Aplosporella prunicola CBS 121.167 v1.0* | Haridas et al. (2020) |
| Bauco1 | Ascomycota | Pezizomycotina | Dothideomycetes | Capnodiales | Teratosphaeriaceae | *Baudoinia compniacensis UAMH 10762 (4089826) v1.0* | Ohm et al. (2012) |
| Aurpu_var_mel1 | Ascomycota | Pezizomycotina | Dothideomycetes | Dothideales | Dothioraceae | *Aureobasidium pullulans var. melanogenum CBS 110374* | Gostincar et al. (2014) |
| Hyspu1_1 | Ascomycota | Pezizomycotina | Dothideomycetes | Hysteriales | Hysteriaceae | *Hysterium pulicare* | Ohm et al. (2012) |
| Micmi1 | Ascomycota | Pezizomycotina | Dothideomycetes | Microthyriales | Microthyriaceae | *Microthyrium microscopicum CBS 115976 v1.0* | Haridas et al. (2020) |
| Elsamp1 | Ascomycota | Pezizomycotina | Dothideomycetes | Myriangiales | Elsinoaceae | *Elsinoe ampelina CECT 20119 v1.0* | Haridas et al. (2020) |
| Lopmy1 | Ascomycota | Pezizomycotina | Dothideomycetes | Mytilinidiales | Mytilinidiaceae | *Lophium mytilinum CBS 269.34 v1.0* | Haridas et al. (2020) |
| Aaoar1 | Ascomycota | Pezizomycotina | Dothideomycetes | Pleosporales | Dacampiaceae | *Aaosphaeria arxii CBS 175.79 v1.0* | Haridas et al. (2020) |
| Tryvi1 | Ascomycota | Pezizomycotina | Dothideomycetes | Trypetheliales | Trypetheliaceae | *Trypethelium eluteriae v1.0* | Haridas et al. (2020) |
| Claca1 | Ascomycota | Pezizomycotina | Eurotiomycetes | Chaetothyriales | Herpotrichiellaceae | *Cladophialophora carrionii CBS 160.54* | Teixeira et al. (2017) |
| Aspacu1 | Ascomycota | Pezizomycotina | Eurotiomycetes | Eurotiales | Aspergillaceae | *Aspergillus aculeatinus CBS 121060 v1.0* | Vesth et al. (2018) |
| Artbe1 | Ascomycota | Pezizomycotina | Eurotiomycetes | Onygenales | Arthrodermataceae | *Arthroderma benhamiae CBS 112371* | Burmester et al. (2011) |
| EndpusZ1 | Ascomycota | Pezizomycotina | Eurotiomycetes | Verrucariales | Verrucariaceae | *Endocarpon pusillum Z07020* | Wang et al. (2014) |
| Pseel1 | Ascomycota | Pezizomycotina | Incertae_sedis | Triblidiales | Triblidiaceae | *Pseudographis elatina* | **Unpublished (1KGF)** |
| Clagr3 | Ascomycota | Pezizomycotina | Lecanoromycetes | Lecanorales | Cladoniaceae | *Cladonia grayi Cgr/DA2myc/ss v2.0* | Armaleo et al. (2019) |
| Lobpul1 | Ascomycota | Pezizomycotina | Lecanoromycetes | Peltigerales | Lobariaceae | *Lobaria pulmonaria Scotland reference genome v1.0* | unpublished (Olafur S. Andrésson) |
| Xanpa2 | Ascomycota | Pezizomycotina | Lecanoromycetes | Teloschistales | Teloschistaceae | *Xanthoria parietina 46-1-SA22 v1.1* | **Unpublished (Paul Dyer)** |
| Acema1 | Ascomycota | Pezizomycotina | Leotiomycetes | Helotiales | Vibrisseaceae | *Acephala macrosclerotiorum EW76-UTF0540 v1.0* | **Unpublished (1KGF)** |
| Amore1 | Ascomycota | Pezizomycotina | Leotiomycetes | Incertae_sedis | Myxotrichaceae | *Amorphotheca resinae v1.0* | Martino et al. (2018) |
| Cocst1 | Ascomycota | Pezizomycotina | Leotiomycetes | Rhytismatales | Rhytismataceae | *Coccomyces strobi CBS 202.91 v1.0* | **Unpublished (1KGF)** |
| Artol1 | Ascomycota | Pezizomycotina | Orbiliomycetes | Orbiliales | Orbiliaceae | *Arthrobotrys oligospora ATCC 24927* | Yang et al. (2011) |
| Aciaci1 | Ascomycota | Pezizomycotina | Pezizomycetes | Pezizales | Amplistromataceae | *Acidothrix acidophila CBS 136259 v1.0* | **Unpublished (1KGF)** |
| Calpu1 | Ascomycota | Pezizomycotina | Sordariomycetes | Calosphaeriales | Calosphaeriaceae | *Calosphaeria pulchella* | **Unpublished (1KGF)** |
| ThoPMI491_1 | Ascomycota | Pezizomycotina | Sordariomycetes | Chaetosphaeriales | Chaetosphaeriaceae | *Thozetella sp. PMI_491 v2.0* | **Unpublished (1KGF)** |
| ConPMI546 | Ascomycota | Pezizomycotina | Sordariomycetes | Coniochaetales | Coniochaetaceae | *Coniochaeta sp. PMI_546 v1.0* | **Unpublished (1KGF)** |
| Crypa2 | Ascomycota | Pezizomycotina | Sordariomycetes | Diaporthales | Cryphonectriaceae | *Cryphonectria parasitica EP155 v2.0* | Crouch et al. (2020) |
| Acrchr1 | Ascomycota | Pezizomycotina | Sordariomycetes | Glomerellales | Plectosphaerellaceae | *Acremonium chrysogenum ATCC 11550* | Terfehr et al. (2014) |
| Beaba1 | Ascomycota | Pezizomycotina | Sordariomycetes | Hypocreales | Cordycipitaceae | *Beauveria bassiana ARSEF 2860* | Xiao et al. (2012) |
| Gaegr1 | Ascomycota | Pezizomycotina | Sordariomycetes | Magnaporthales | Magnaporthaceae | *Gaeumannomyces graminis var. tritici R3-111a-1* | Okagaki et al. (2015) |
| Melti1 | Ascomycota | Pezizomycotina | Sordariomycetes | Melanosporales | Melanosporaceae | *Melanospora tiffanyae F1KG0001 v1.0* | **Unpublished (1KGF)** |
| Corma2 | Ascomycota | Pezizomycotina | Sordariomycetes | Microascales | Halosphaeriaceae | *Corollospora maritima CBS 119819 v2.0* | **Unpublished (1KGF)** |
| Grocl1 | Ascomycota | Pezizomycotina | Sordariomycetes | Ophiostomatales | Ophiostomataceae | *Grosmannia clavigera kw1407* | DiGuistini et al. (2011) |
| Achstr1 | Ascomycota | Pezizomycotina | Sordariomycetes | Sordariales | Chaetomiaceae | *Achaetomium strumarium CBS333.67 v1.0* | **Unpublished (Pierre Gladieux, Francis Martin)** |
| Khuory1 | Ascomycota | Pezizomycotina | Sordariomycetes | Trichosphaeriales | Trichosphaeriaceae | *Khuskia oryzae ATCC 28132 v1.0* | **Unpublished (1KGF)** |
| Antav1 | Ascomycota | Pezizomycotina | Sordariomycetes | Xylariales | Diatrypaceae | *Anthostoma avocetta NRRL 3190 v1.0* | **Unpublished (1KGF)** |
| Trigu1 | Ascomycota | Pezizomycotina | Xylonomycetes | Xylonales | Xylonaceae | *Trinosporium guianense CBS132537 v1.0* | **Unpublished (1KGF)** |
| Neoirr1 | Ascomycota | Taphrinomycotina | Neolectomycetes | Neolectales | Neolectaceae | *Neolecta irregularis DAH-1 v1.0* | Nguyen et al. (2017) |
| Agabi_varbisH97_2 | Basidiomycota | Agaricomycotina | Agaricomycetes | Agaricales | Agaricaceae | *Agaricus bisporus var bisporus (H97) v2.0* | Morin et al. (2012) |
| Fibsp1 | Basidiomycota | Agaricomycotina | Agaricomycetes | Atheliales | Atheliaceae | *Fibulorhizoctonia sp. CBS 109695 v1.0* | Nagy et al. (2016) |
| Aurde3_1 | Basidiomycota | Agaricomycotina | Agaricomycetes | Auriculariales | Auriculariaceae | *Auricularia subglabra v2.0* | Floudas et al. (2012) |
| Boled1 | Basidiomycota | Agaricomycotina | Agaricomycetes | Boletales | Boletaceae | *Boletus edulis v1.0* | **Unpublished (Francis Martin)** |
| Botbo1 | Basidiomycota | Agaricomycotina | Agaricomycetes | Cantharellales | Botryobasidiaceae | *Botryobasidium botryosum v1.0* | Riley et al. (2014) |
| Punst1 | Basidiomycota | Agaricomycotina | Agaricomycetes | Corticiales | Punctulariaceae | *Punctularia strigosozonata v1.0* | Floudas et al. (2012) |
| Sphst1 | Basidiomycota | Agaricomycotina | Agaricomycetes | Geastrales | Sphaerobolaceae | *Sphaerobolus stellatus v1.0* | Kohler et al. (2015) |
| Glotr1_1 | Basidiomycota | Agaricomycotina | Agaricomycetes | Gloeophyllales | Gloeophyllaceae | *Gloeophyllum trabeum v1.0* | Floudas et al. (2012) |
| Gaumor1_1 | Basidiomycota | Agaricomycotina | Agaricomycetes | Gomphales | Gautieriaceae | *Gautieria morchelliformis GMNE.BST v1.0* | **Unpublished (1KGF)** |
| Fomme1 | Basidiomycota | Agaricomycotina | Agaricomycetes | Hymenochaetales | Hymenochaetaceae | *Fomitiporia mediterranea v1.0* | Floudas et al. (2012) |
| Jaaar1 | Basidiomycota | Agaricomycotina | Agaricomycetes | Jaapiales | Jaapiaceae | *Jaapia argillacea v1.0* | Riley et al. (2014) |
| Mutel1 | Basidiomycota | Agaricomycotina | Agaricomycetes | Phallales | Phallaceae | *Mutinus elegans ME.BST v1.0* | **Unpublished (1KGF)** |
| Abobie1 | Basidiomycota | Agaricomycotina | Agaricomycetes | Polyporales | Meruliaceae | *Abortiporus biennis CIRM-BRFM1778 v1.0* | **Unpublished (Marie-Noëlle Rosso)** |
| Amycha1 | Basidiomycota | Agaricomycotina | Agaricomycetes | Russulales | Amylostereaceae | *Amylostereum chailletii DWAch2 v1.0* | **Unpublished (Francis Martin)** |
| Pirin1 | Basidiomycota | Agaricomycotina | Agaricomycetes | Sebacinales | Sebacinaceae | *Piriformospora indica DSM 11827 from MPI* | Zuccaro et al. (2011) |
| Thega1 | Basidiomycota | Agaricomycotina | Agaricomycetes | Thelephorales | Thelephoraceae | *Thelephora ganbajun P2 v1.0* | **Unpublished (1KGF)** |
| Sisni1 | Basidiomycota | Agaricomycotina | Agaricomycetes | Trechisporales | Trechisporaceae | *Sistotremastrum niveocremeum HHB9708 ss-1 1.0* | Nagy et al. (2016) |
| Calco1 | Basidiomycota | Agaricomycotina | Dacrymycetes | Dacrymycetales | Dacrymycetaceae | *Calocera cornea v1.0* | Nagy et al. (2016) |
| Walse1 | Basidiomycota | Agaricomycotina | Wallemiomycetes | Wallemiales | Wallemiaceae | *Wallemia mellicola v1.0* | Padamsee et al. (2012) |
| Atrsp2 | Basidiomycota | Pucciniomycotina | Atractiellomycetes | Atractiellales | Incertae_sedis | *Atractiellales rhizophila v2.0* | **Unpublished (Francis Martin, Greg Bonito)** |
| Croqu1 | Basidiomycota | Pucciniomycotina | Pucciniomycetes | Pucciniales | Cronartiaceae | *Cronartium quercuum f. sp. fusiforme G11 v1.0* | Pendleton et al. (2014) |

**Supplementary Table 4.** Number of gene models with evidence for HGT associated to taxonomy categories from the NCBI RefSeq database.

| **NCBI taxonomy** | *B. meristosporus* CBS 931.73 | *B.meristosporus* B9252 | *B. heterosporus* B8290 |
| --- | --- | --- | --- |
| a-proteobacteria | 49 | 33 | 21 |
| b-proteobacteria | 39 | 31 | 21 |
| d-proteobacteria | 56 | 40 | 23 |
| e-proteobacteria | 6 | 1 | 2 |
| g-proteobacteria | 82 | 54 | 27 |
| proteobacteria | 2 | 1 | 1 |
| firmicutes | 112 | 87 | 54 |
| actinobacteria | 2 | 4 | 1 |
| high GC Gram+ | 125 | 63 | 39 |
| enterobacteria | 13 | 10 | 2 |
| planctomycetes | 18 | 6 | 2 |
| CFB group bacteria | 79 | 60 | 33 |
| bacteriodetes | 91 | 28 | 12 |
| verrucomicrobia | 10 | 8 | 5 |
| fusobacteria | NA | NA | 2 |
| cyanobacteria | 89 | 51 | 42 |
| chlamydias | NA | NA | 1 |
| mycoplasmas | 1 | NA | 1 |
| aquificales | 1 | NA | 1 |
| bacteria | 28 | 21 | 6 |
| euryarchaeotes | 6 | 4 | 4 |
| archaea | 1 | 1 | 1 |
| **Total** | **810** | **503** | **301** |

**Supplementary Table 5.** Number of GO terms and SM categories annotated from gene models with evidence for HGT in *B. meristosporus* CBS 931.73

| **Gene Ontology term** | **Number of gene models** |
| --- | --- |
| No GO | 659 |
| No InterPro | 140 |
| GO:oxidation-reduction process | 59 |
| GO:protein binding | 34 |
| GO:phosphorelay sensor kinase activity | 32 |
| GO:N-acetyltransferase activity | 28 |
| GO:catalytic activity | 26 |
| GO:extracellular space | 25 |
| GO:hydrolase activity | 24 |
| GO:oxidoreductase activity | 23 |
| GO:hydrolase activity, hydrolyzing O-glycosyl compounds | 19 |
| **NRPS** | 18 |
| GO:amino acid transport | 13 |
| GO:transport | 13 |
| GO:methyltransferase activity | 11 |
| GO:proteolysis | 10 |
| GO:ATP binding | 7 |
| GO:FMN binding | 6 |
| GO:hydrolase activity, acting on carbon-nitrogen (but not peptide) bonds | 6 |
| GO:DNA binding | 5 |
| **NRPS-Like** | 4 |
| GO:carbon-nitrogen ligase activity, with glutamine as amido-N-donor | 4 |
| GO:chitinase activity | 4 |
| GO:catalase activity | 3 |
| GO:cellular amino acid metabolic process | 3 |
| GO:FAD binding | 3 |
| GO:nucleic acid binding | 3 |
| GO:oxidoreductase activity, acting on other nitrogenous compounds as donors | 3 |
| GO :transferase activity, transferring acyl groups other than amino-acyl groups | 3 |
| GO:aminoacyl-tRNA editing activity | 2 |
| GO:coproporphyrinogen oxidase activity | 2 |
| GO:nucleotide binding | 2 |
| GO:nutrient reservoir activity | 2 |
| GO:phosphogluconate dehydrogenase (decarboxylating) activity | 2 |
| GO:response to oxidative stress | 2 |
| GO:serine-type endopeptidase activity | 2 |
| GO:transmembrane transport | 2 |
| GO:tryptophan synthase activity | 2 |
| GO:zinc ion binding | 2 |
| **PKS** | 2 |
| **Terpene** | 2 |
| GO:cation transmembrane transporter activity | 1 |
| GO:copper ion binding | 1 |
| GO:dihydroorotate dehydrogenase activity | 1 |
| GO:hydrolase activity, acting on glycosyl bonds | 1 |
| GO:integral component of membrane | 1 |
| GO:iron ion binding | 1 |
| GO:lipid metabolic process | 1 |
| GO:lyase activity | 1 |
| GO:MAP kinase activity | 1 |
| GO:nucleobase-containing compound metabolic process | 1 |
| GO:orotate phosphoribosyltransferase activity | 1 |
| GO:oxidoreductase activity, acting on CH-OH group of donors | 1 |
| GO:polysaccharide catabolic process | 1 |
| GO:protein tyrosine phosphatase activity | 1 |
| GO:protein-chromophore linkage | 1 |
| GO:pyridoxal phosphate binding | 1 |
| GO:regulation of transcription, DNA-templated | 1 |
| GO:RNA binding | 1 |
| GO:structural molecule activity | 1 |
| GO:ubiquitin-dependent protein catabolic process | 1 |
