## Supplementary Figures for "Phylogenomic analyses of non-Dikarya fungi supports horizontal gene transfer driving diversification of secondary metabolism in the amphibian gastrointestinal symbiont, *Basidiobolus*"

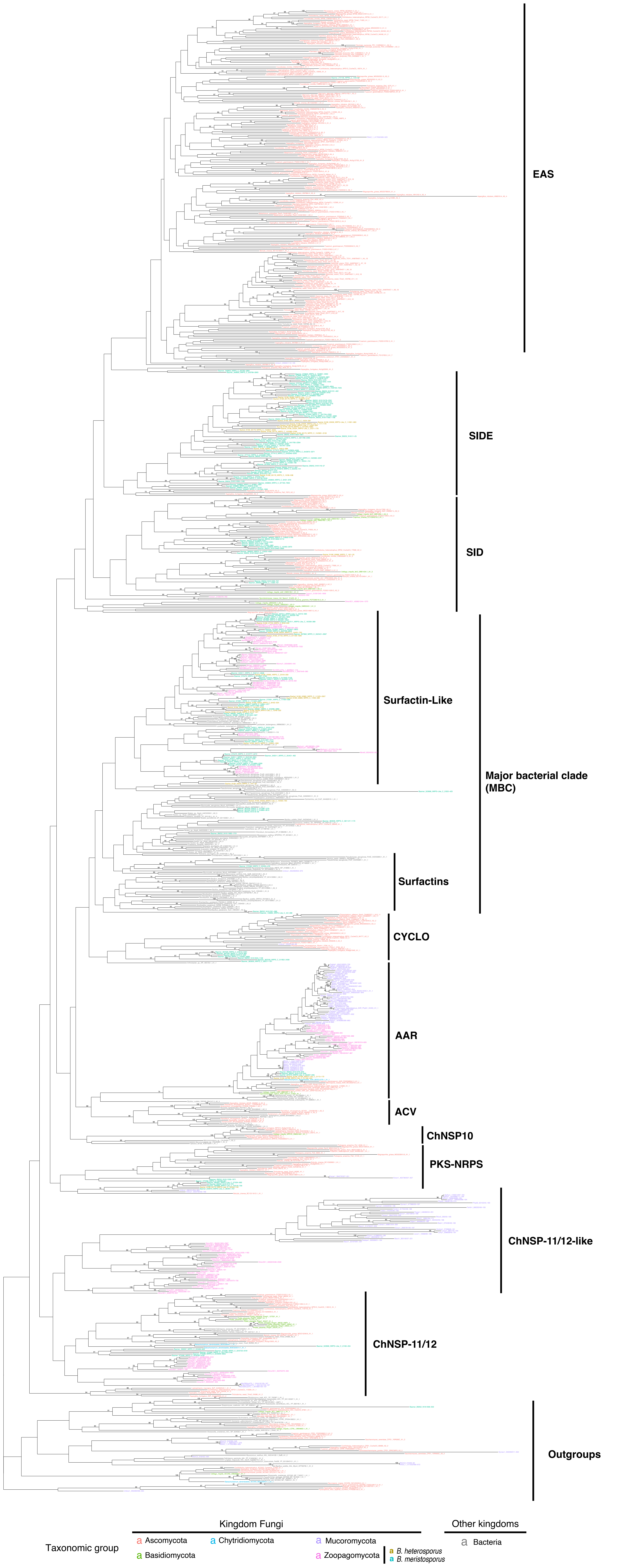


Supplementary Figure 1. Maximum likelihood tree reconstruction using NRPS A-domains from Zoopagomycota and Mucoromycota genomes. The maximum likelihood phylogenetic was reconstructed using the A-domains from Bushley et al. (2010) and the A-domains predicted for all genomes used in this report. Numbers above each branch represent bootstrap values after 1,000 replicates when support is greater than 70%. Cluster names represent the groups assigned to each A-domain clade.

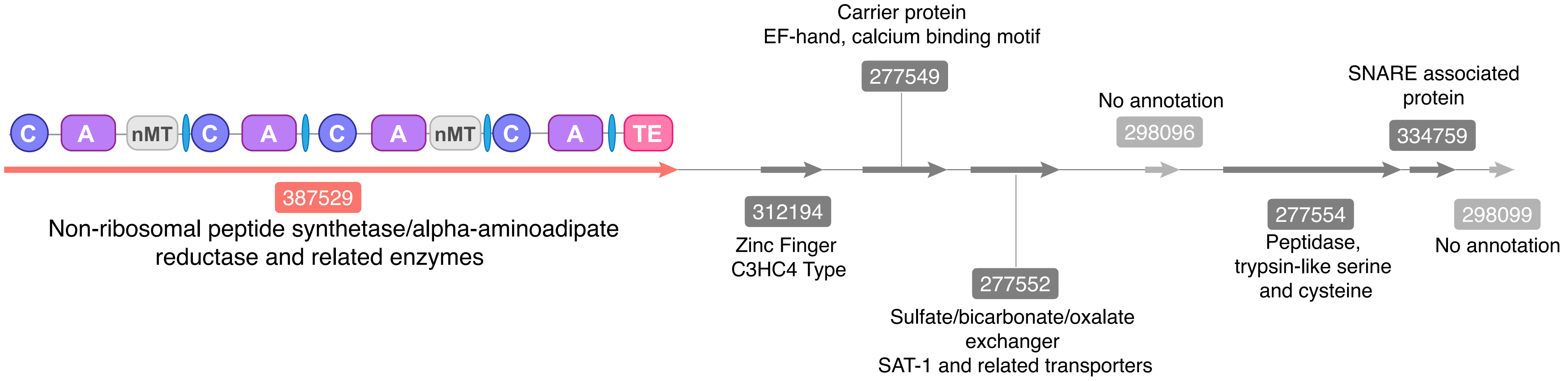


**NRPS Cluster 5 (*B. meristosporus* CBS 931.73)**

Supplementary Figure 2. Graphic representation of the Cluster 5 (NRPS) of *B. meristosporus* CBS 931.73. The SM gene model associated to Cluster 5 ( 387529; NRPS) contains 4 A-domains clustered within the CYCLO clade of the NRPS phylogenetic tree (Figure 3). This clade contains the A-domains of *Tolypocladium inflatum* cyclosporin NRPS gene. Domains of the NRPS protein are illustrated above the arrow representing the gene. Dark purple: Condensation; Light Purple: Adenylation; Grey: n-Methylation; Pink: Thioesterase. The arrows in grey represent the additional gene models predicted to be part of Cluster 5.

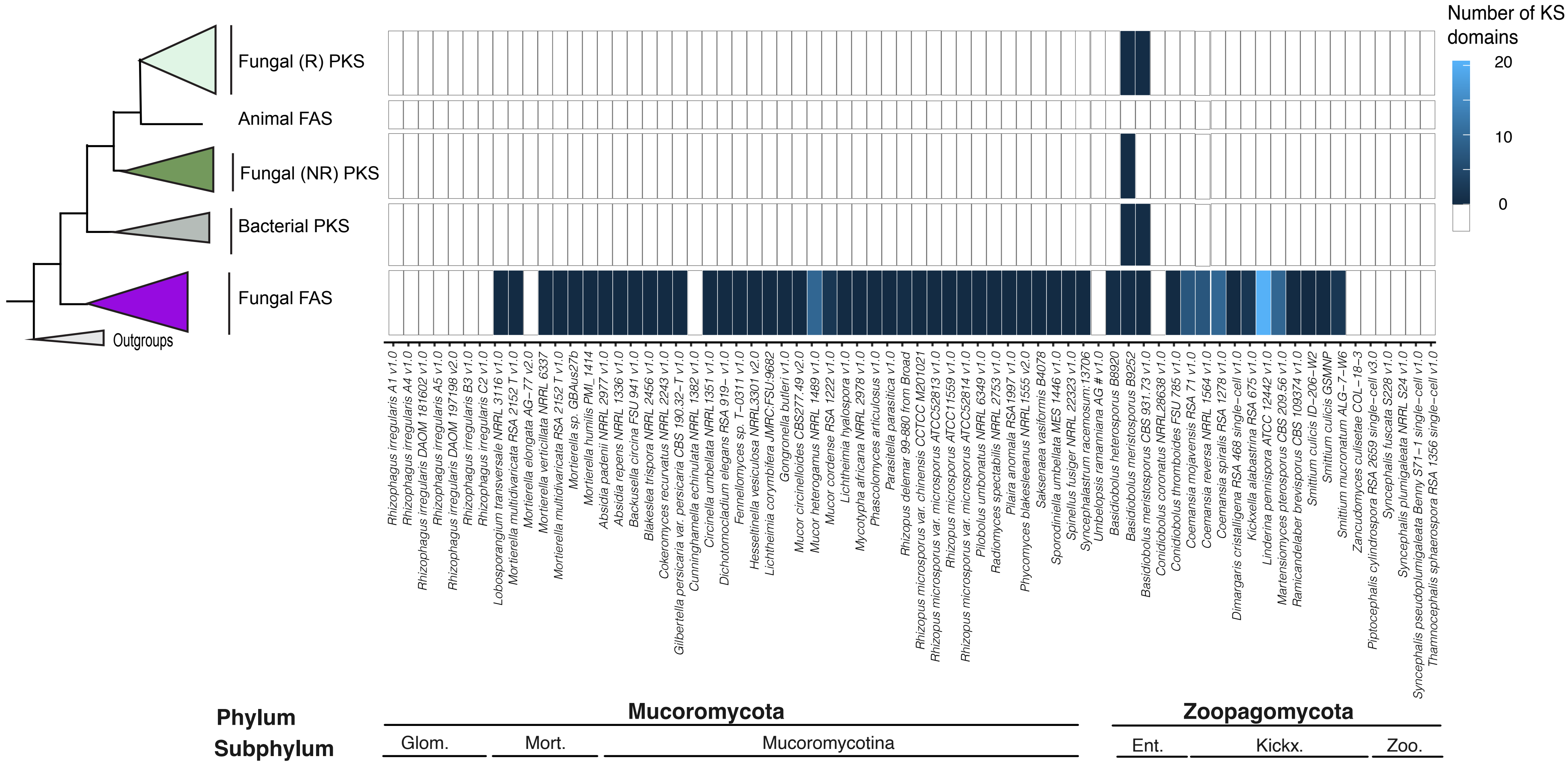


Supplementary Figure 3. Phylogenetic sources and abundance of KS-domains from PKS predicted gene models for Zoopagomycota and Mucoromycota species. The heatmap (right) represents the abundances KS-domains predicted for each domain clustered within each clade. Glom.: Glomeromycotina. Mort.: Morteriellomycotina. Ent.: Entomophtoromycotina. Kickx.: Kickxellomycotina. Zoo.: Zoopagomycotina

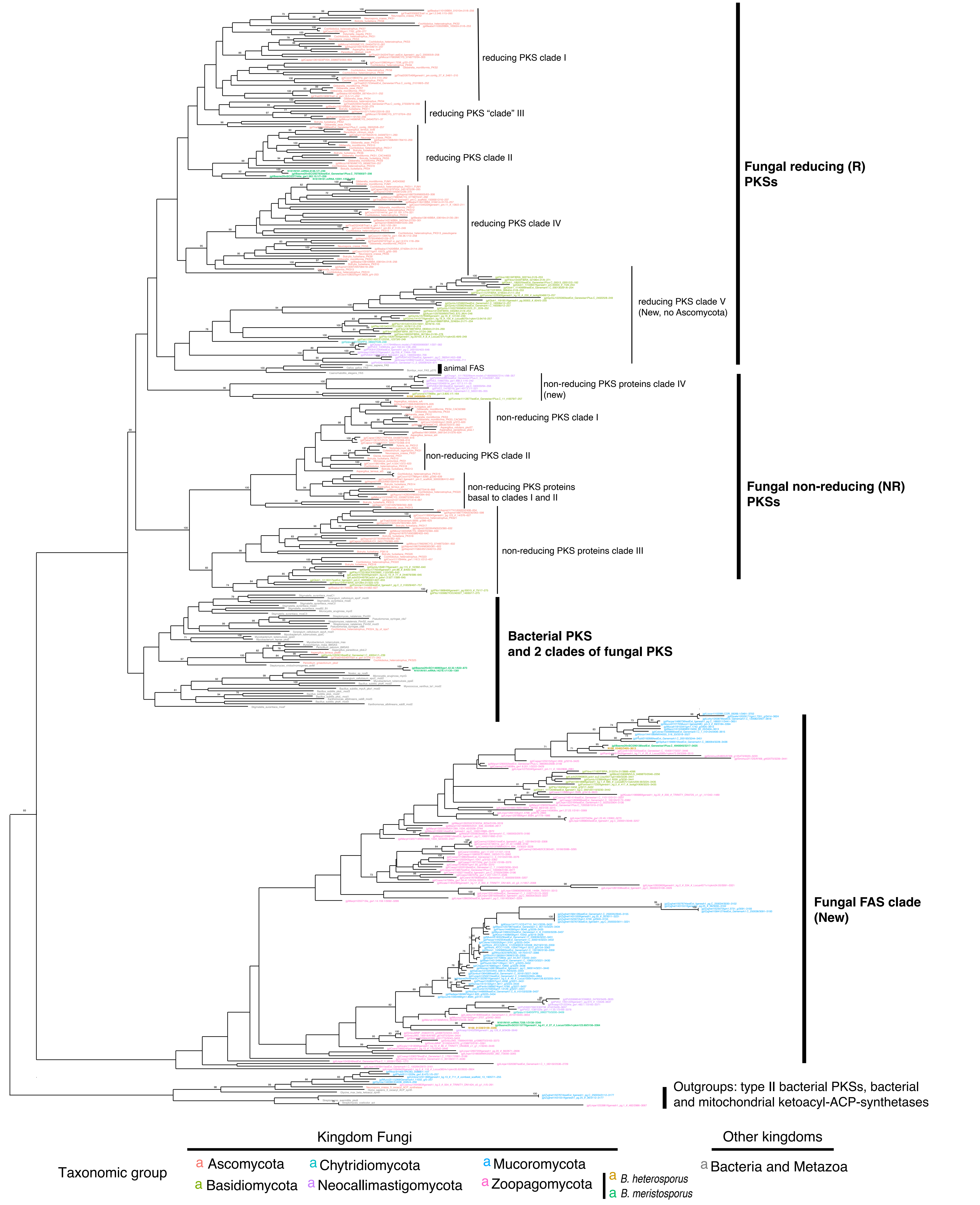


Supplementary Figure 4. Maximum likelihood tree reconstruction using PKS KS-domains from Zoopagomycota and Mucoromycota genomes. The maximum likelihood phylogenetic was reconstructed using the KS-domains from Kroker et al. (2013) and the KS domains predicted for all genomes used in this report. Numbers above each branch represent bootstrap values after 1,000 replicates when support is greater than 70%. Cluster names represent the groups assigned to each KS-domain clade.

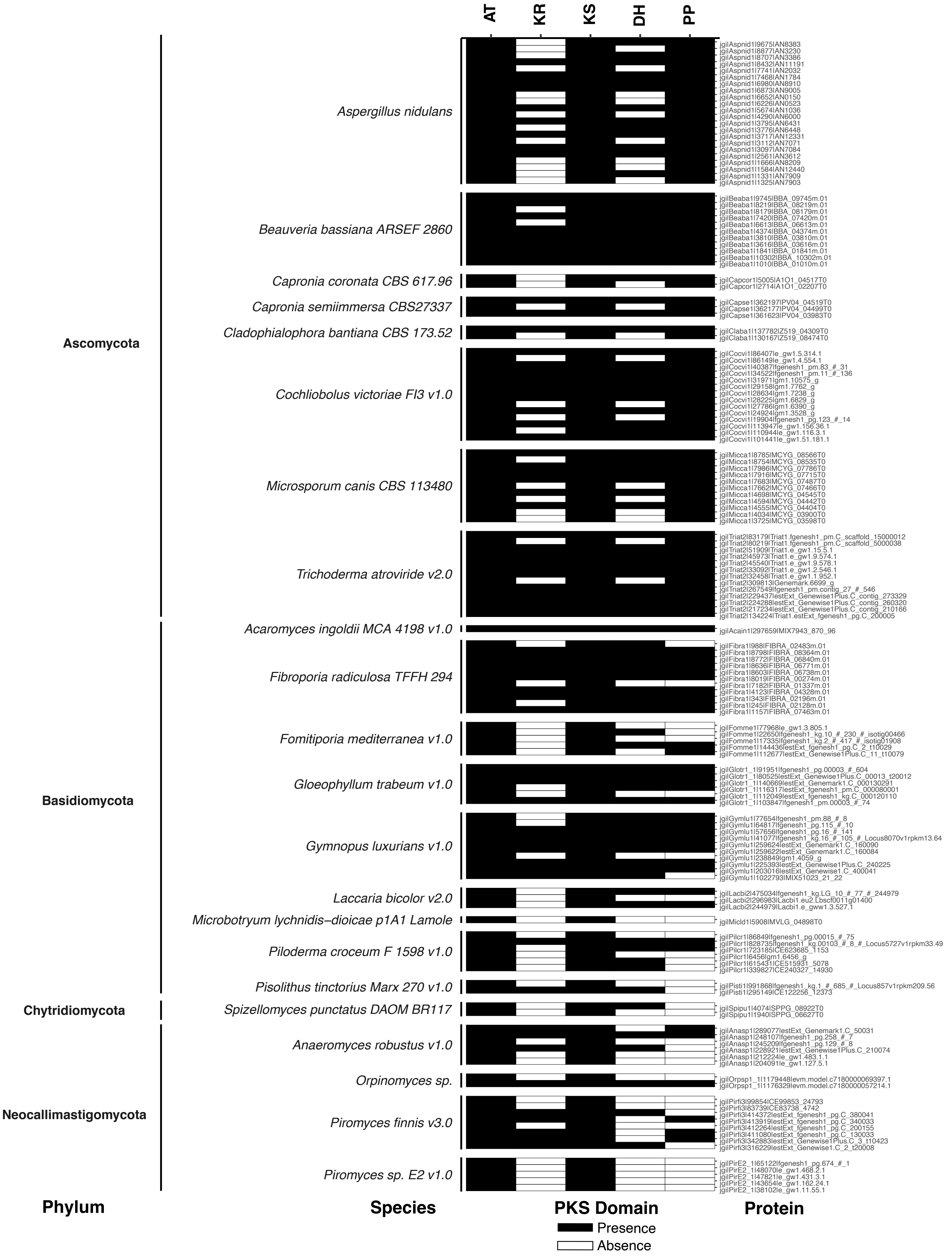


Supplementary Figure 5. Presence and absence of domains characteristic of PKS core gene models in non-zygomycete Fungal species. Shaded colors indicate presence.

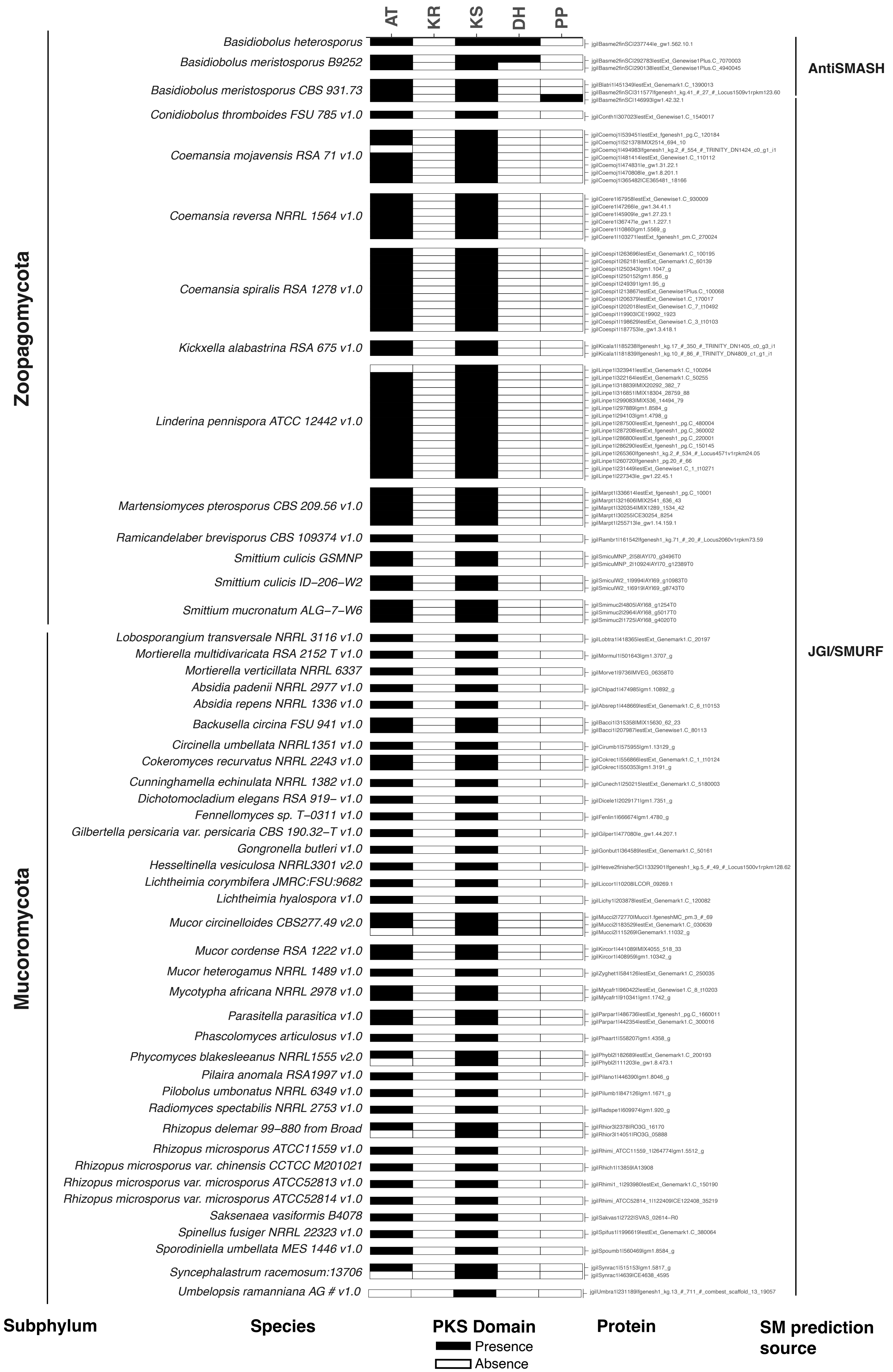


Supplementary Figure 6. Presence and absence of domains characteristic of PKS core gene models in zygomycete species. Shaded colors indicate presence.

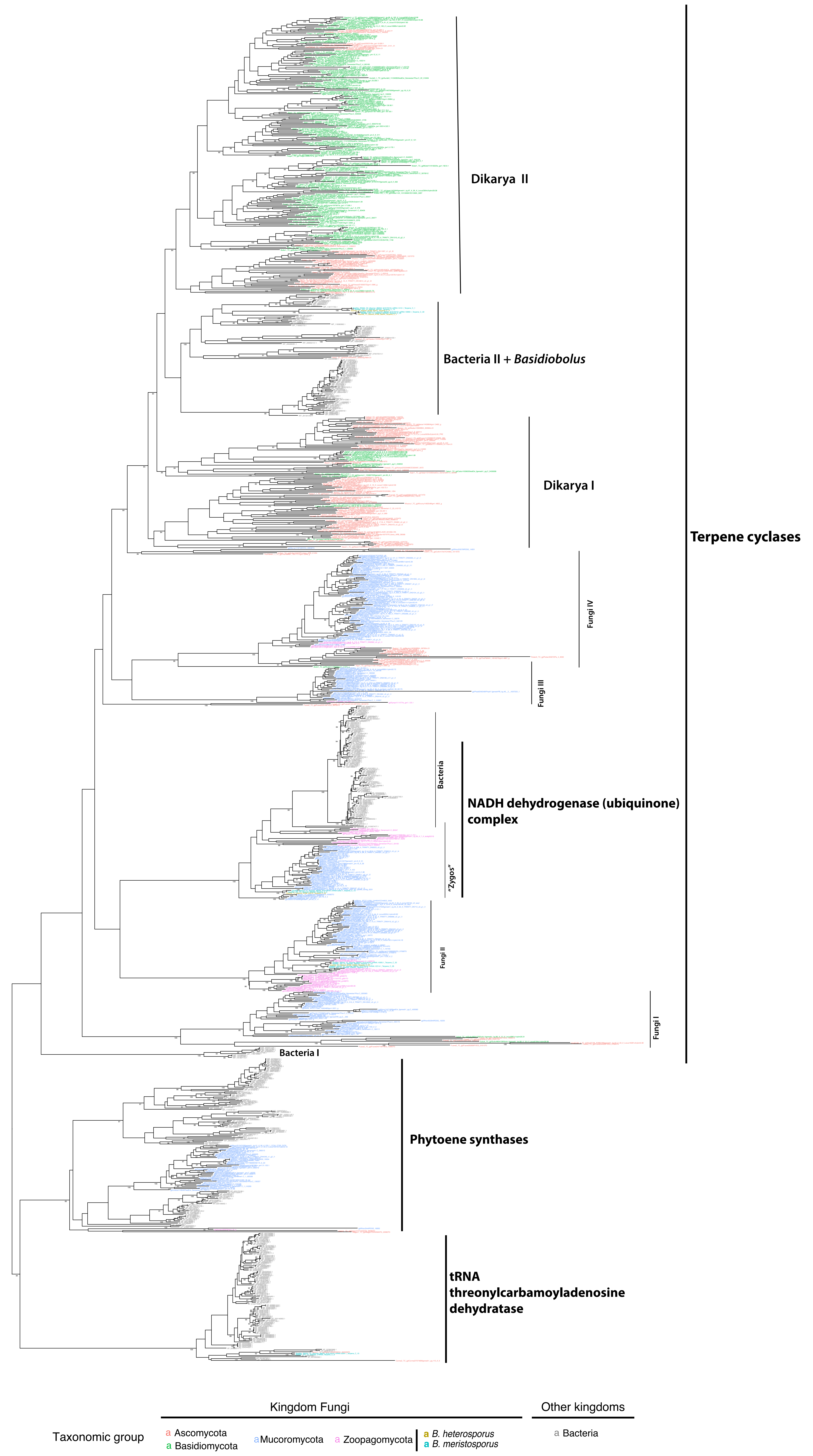


Supplementary Figure 7. Maximum likelihood tree reconstruction using the predicted terpene cyclase (TC) core genes from Zoopagomycota and Mucoromycota genomes. The maximum likelihood phylogenetic was reconstructed using the predicted TC domains for the genomes of interest. Numbers above each branch represent bootstrap values after 1,000 replicates when support is greater than 70%.

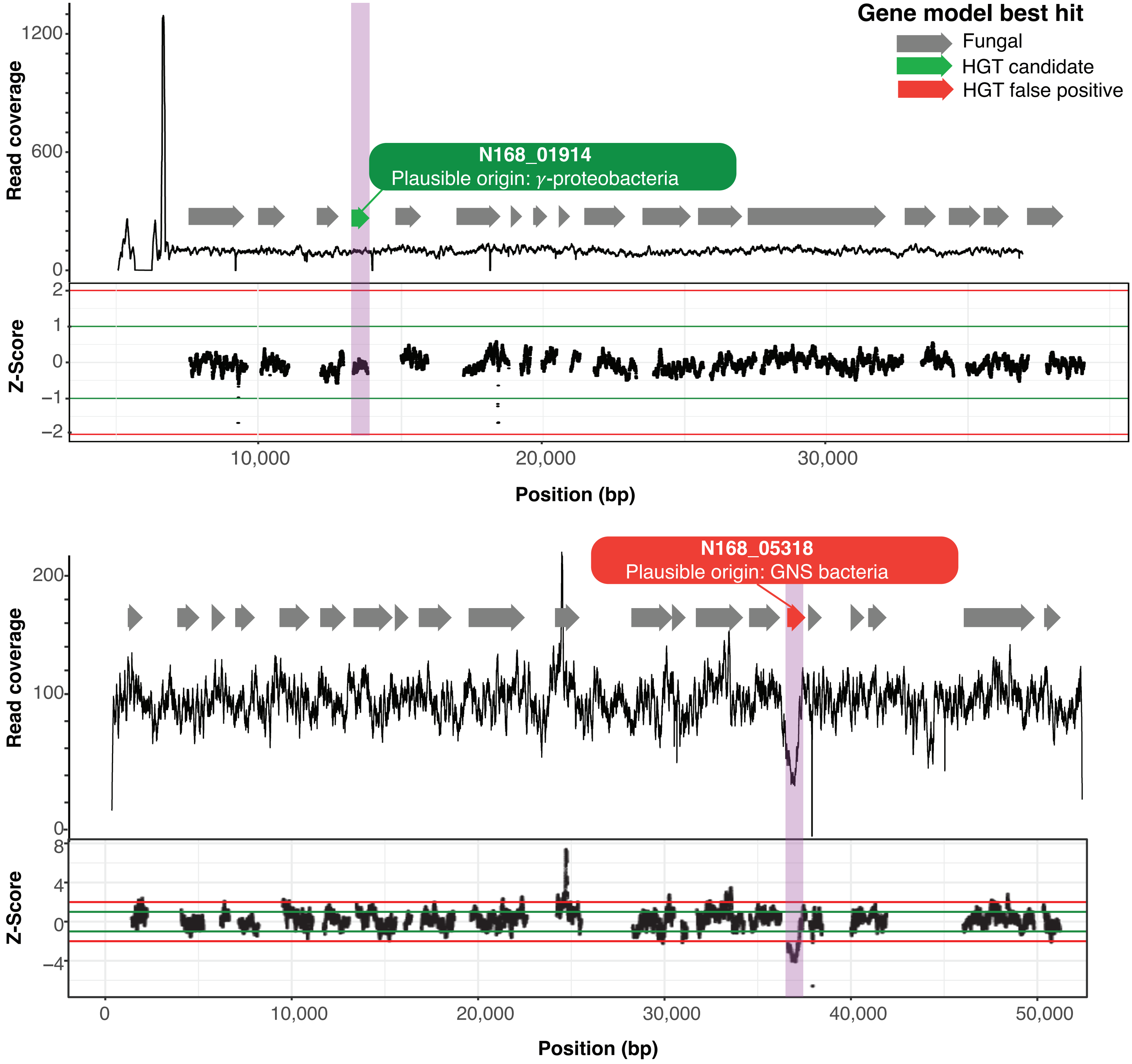


Supplementary Figure 8. Illustration of HGT assay to detect HGT candidates. Figure shows the read coverage and z-score per position of scaffold jcf7180000797043 from *B. heterosporus* B8920. Z-score represents the number of standard deviations from the mean coverage per genic position; any gene model with a z-score greater than 2 or lower than -2 is removed from the analysis. BLAST results show that N168_01914 best hit is to alpha-proteobacteria from RefSeq. Coverage and z-scores for gene model N168_01914 do not show significant deviation of coverage (purple rectangle), making N168_01914 a plausive candidate for HGT origin.

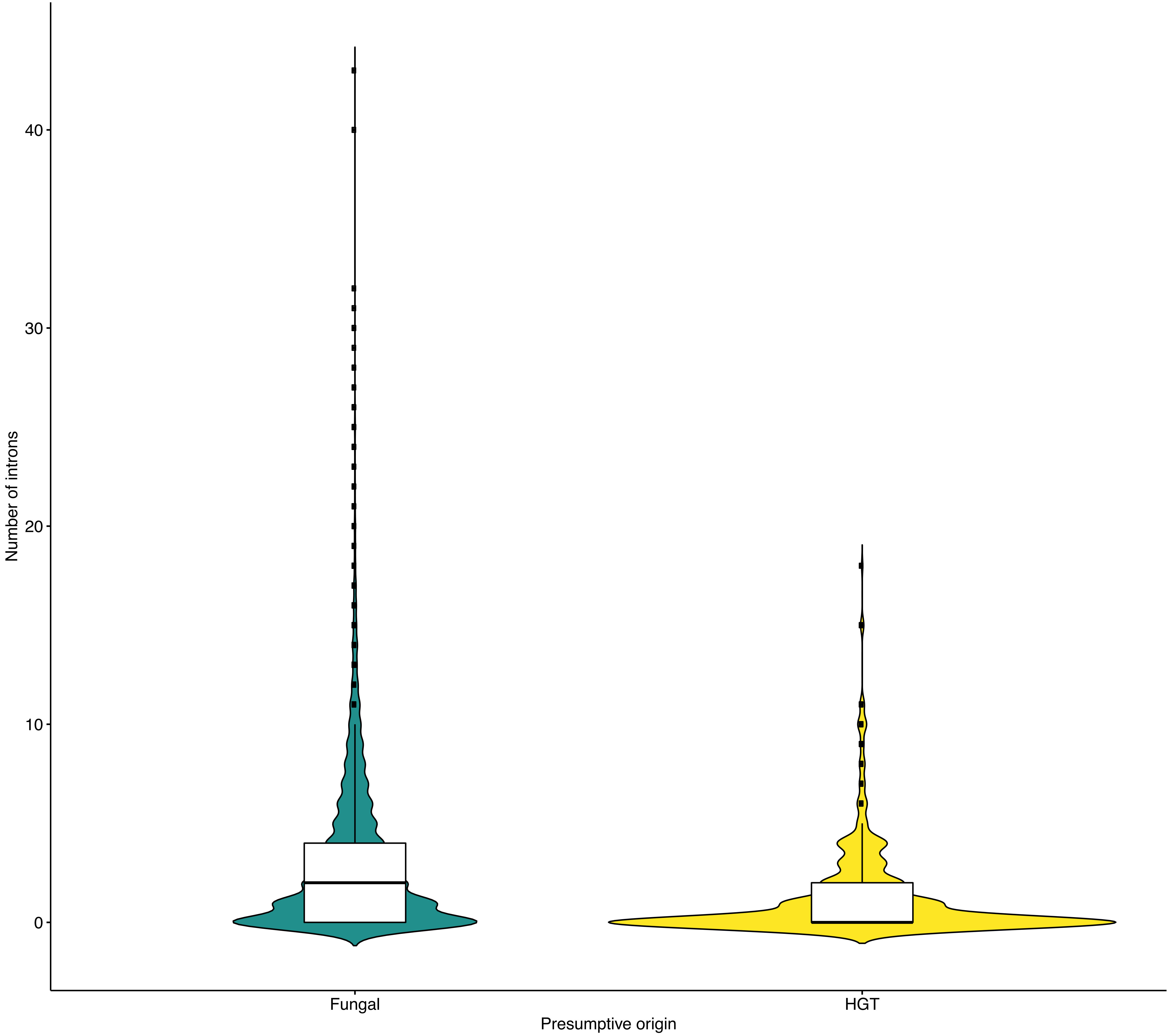


Supplementary Figure 9. Distribution and number of introns for the genes from the HGT assay idenfied in *Basidiobolus meristosporus* CBS 931.73. Distribution of quantiles is represented by the boxplots, where bold lines represent the median number of introns per genic origin. The violin plot represents the raw distribution of the data. Colors represent origin. A higher number of genes with no introns are found in genes of presumptive HGT origin than in fungal genes.

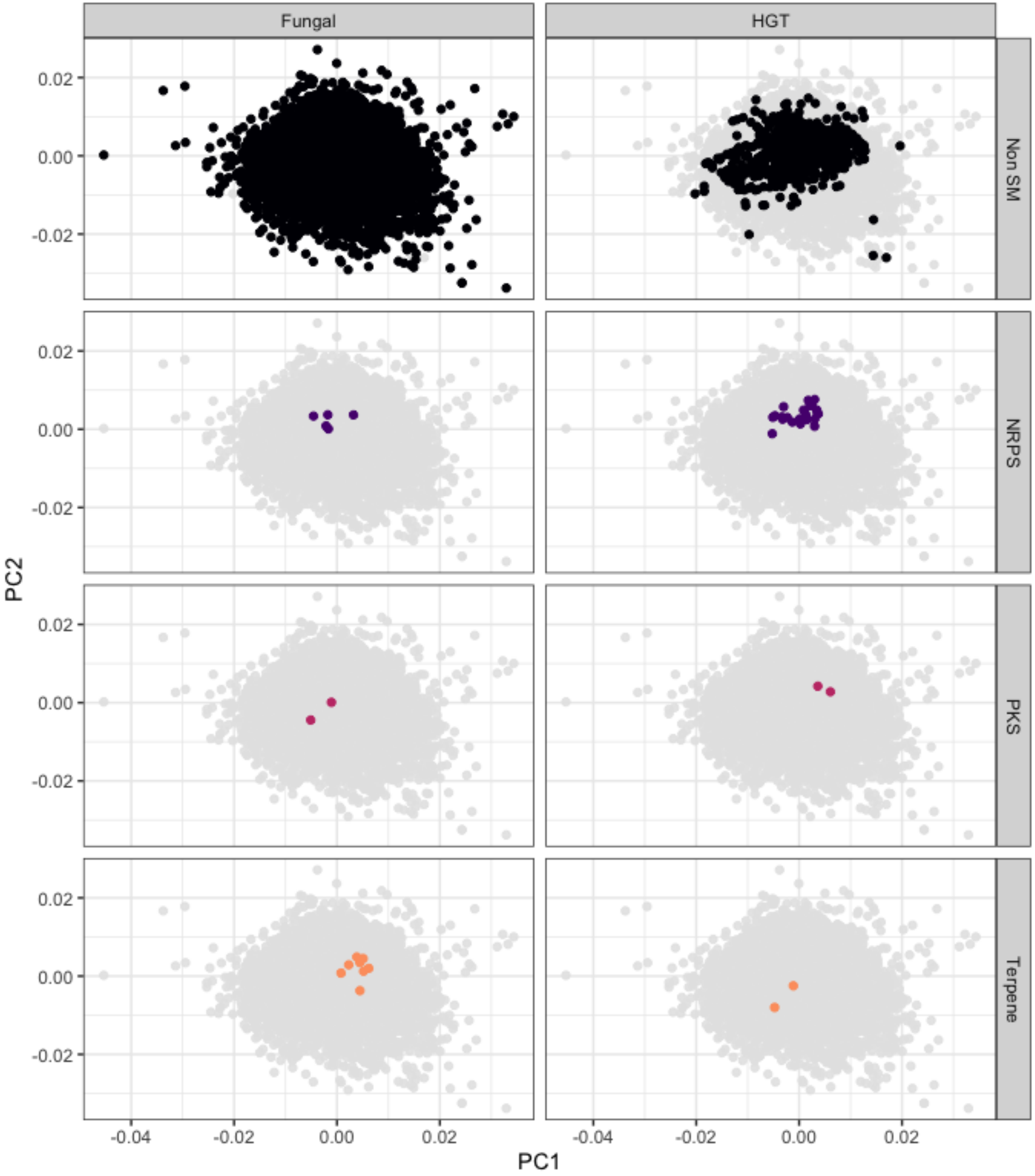


Supplementary Figure 10. Principal components analysis of 5-meres for the genes predicted from *Basidiobolus meristosporus* CBS 931.73 from the HGT assay. Rows represent secondary metabolite classes and columns putative origin. The two principal components with the highest variance contribution (PC1, 14% and PC2, 8%) are shown. Points represent the genes from the HGT analysis. Colors represent secondary metabolite classes.

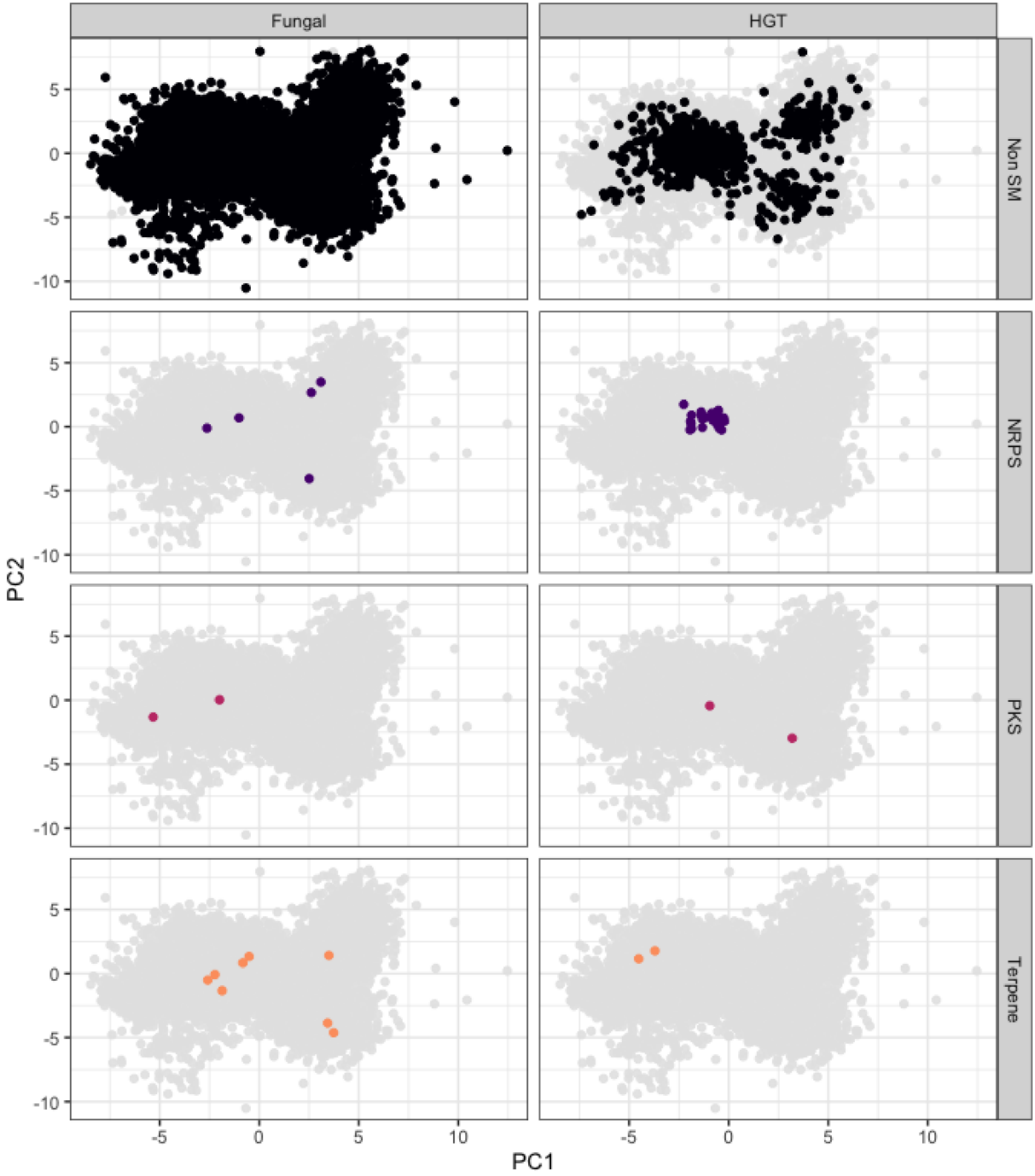


Supplementary Figure 11. Principal components analysis of codon usage for the genes predicted from *Basidiobolus meristosporus* CBS 931.73 from the HGT assay. Rows represent secondary metabolite classes and columns putative origin. The two principal components with the highest variance contribution (PC1, 3% and PC2, 2%) are shown. Points represent the genes from the HGT analysis. Colors represent secondary metabolite classes.

*Basidiobolus meristosporus* CBS 931.73

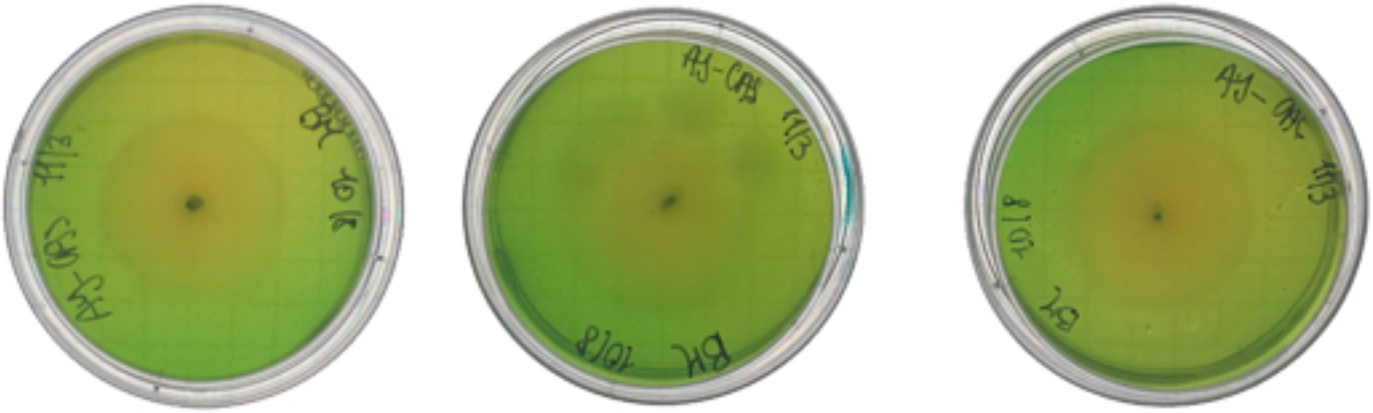

*Cladosporium sp.* PE-07

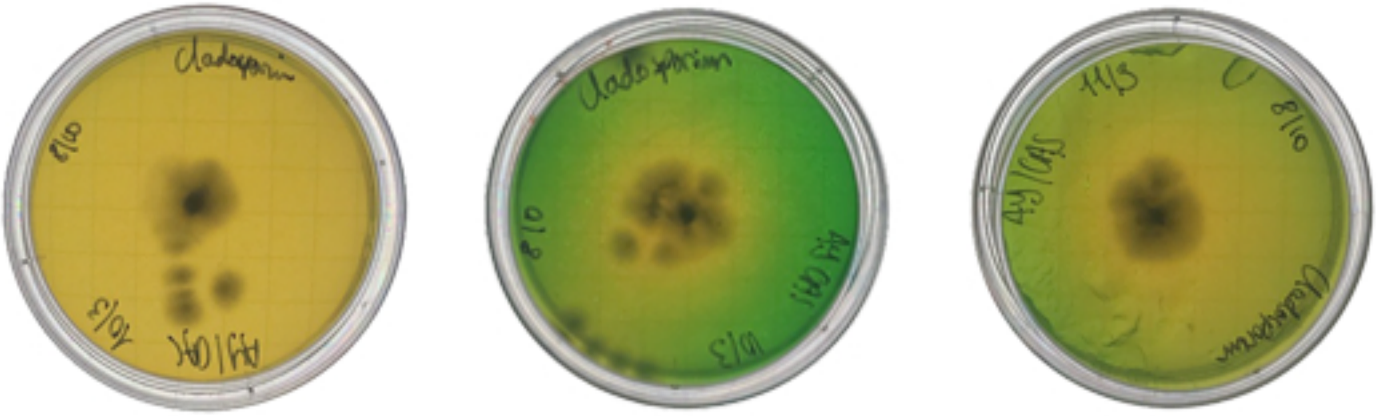

*Conidiobolus thromboides* FSU 785

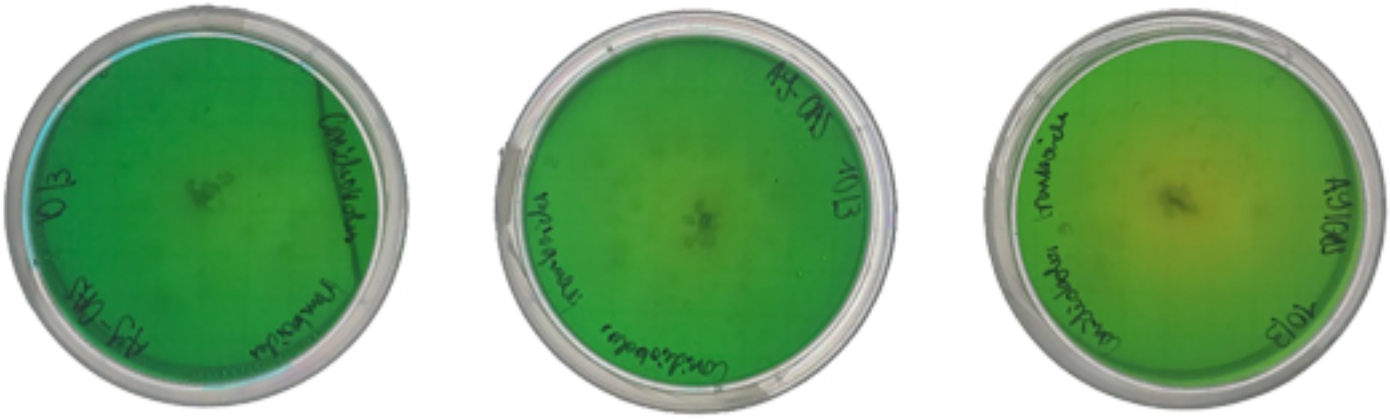

Empty plates (Negative control)

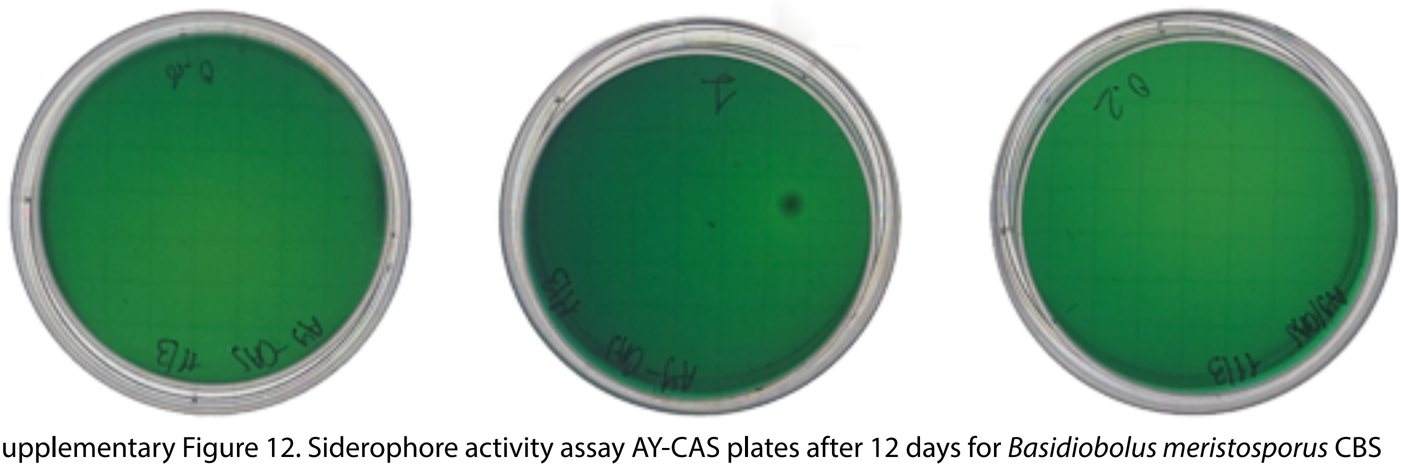

Supplementary Figure 12. Siderophore activity assay AY-CAS plates after 12 days for *Basidiobolus meristosporus* CBS 931.73, *Cladosporium sp*. PE-07, *Conidiobolus thromboides* FSU-785, and an empty plate as negative control. The AY-CAS starts with the greenish-blue color as shown by the empty plates, and turns yellow as siderophores remove the iron from the CAS assay complex.
